# Comparison of Culture Systems for Mouse Living Myocardial Slices in Cardiac Fibrosis Research

**DOI:** 10.64898/2026.08.19.745687

**Authors:** Nora Kopse, Gino A. Bonazza, Andrea Laimbacher, Astrid Hofman, Oliver Distler, Przemyslaw Blyszczuk, Gabriela Kania

**Affiliations:** Center of Experimental Rheumatology, Department of Rheumatology, University Hospital Zurich, University of Zurich, 8952 Schlieren, Switzerland; Department of Clinical Immunology, Jagiellonian University Medical College, Cracow, Poland

**Keywords:** living myocardial slices, cardiac fibrosis, biomimetic system

## Abstract

Living myocardial slices (LMS) are a highly relevant ex vivo model for investigating cardiac physiology and disease, as they preserve the native three-dimensional architecture, cellular diversity, and extracellular matrix of the heart. In addition, LMS enable longitudinal functional and molecular analyses. In this study, we established and compared two LMS culture approaches: an air-liquid interface system and a biomimetic culture system. We further examined how different slicing techniques affect tissue quality and longevity within the biomimetic setup.

To develop a fibrosis model, LMS were stimulated with transforming growth factor-β1 (TGF-β1) and/or exposed to increased mechanical load. Tissue viability was assessed using LIVE/DEAD staining and the MTT assay, while cytotoxicity was evaluated with the LDH-Glo™ Cytotoxicity assay. Contractile function was measured, and fibrotic remodelling was analysed using RT-qPCR, ELISA, and immunohistochemistry.

Our results demonstrate that LMS cultured in the biomimetic system exhibit superior viability, structural integrity, and functional performance compared with those maintained at the air-liquid interface. Mouse LMS could be stably cultured for up to one week in the biomimetic system. Importantly, sample preparation, particularly the slicing method, had a significant impact on tissue quality and culture duration. While TGF-β1 stimulation alone did not consistently induce fibrosis, combining TGF-β1 treatment with increased mechanical load led to more pronounced fibrotic remodelling in LMS. These findings highlight the importance of biomechanical cues in modelling cardiac fibrosis ex vivo and support the biomimetic system as a robust platform for functional and disease-relevant studies.

**Highlights:**

- Biomimetic culture preserves viability, structure, and contractile function of mouse LMS better than air-quid interface culture.
- Sample preparation, specifically the slicing technique, critically determines LMS quality and culture longevity.
- Mouse LMS can be maintained for up to six days in a biomimetic system, enabling longitudinal functional analyses.
- Increased mechanical load and culture duration, not TGF-β1, drove fibrotic changes in LMS.

**Graphical abstract:** 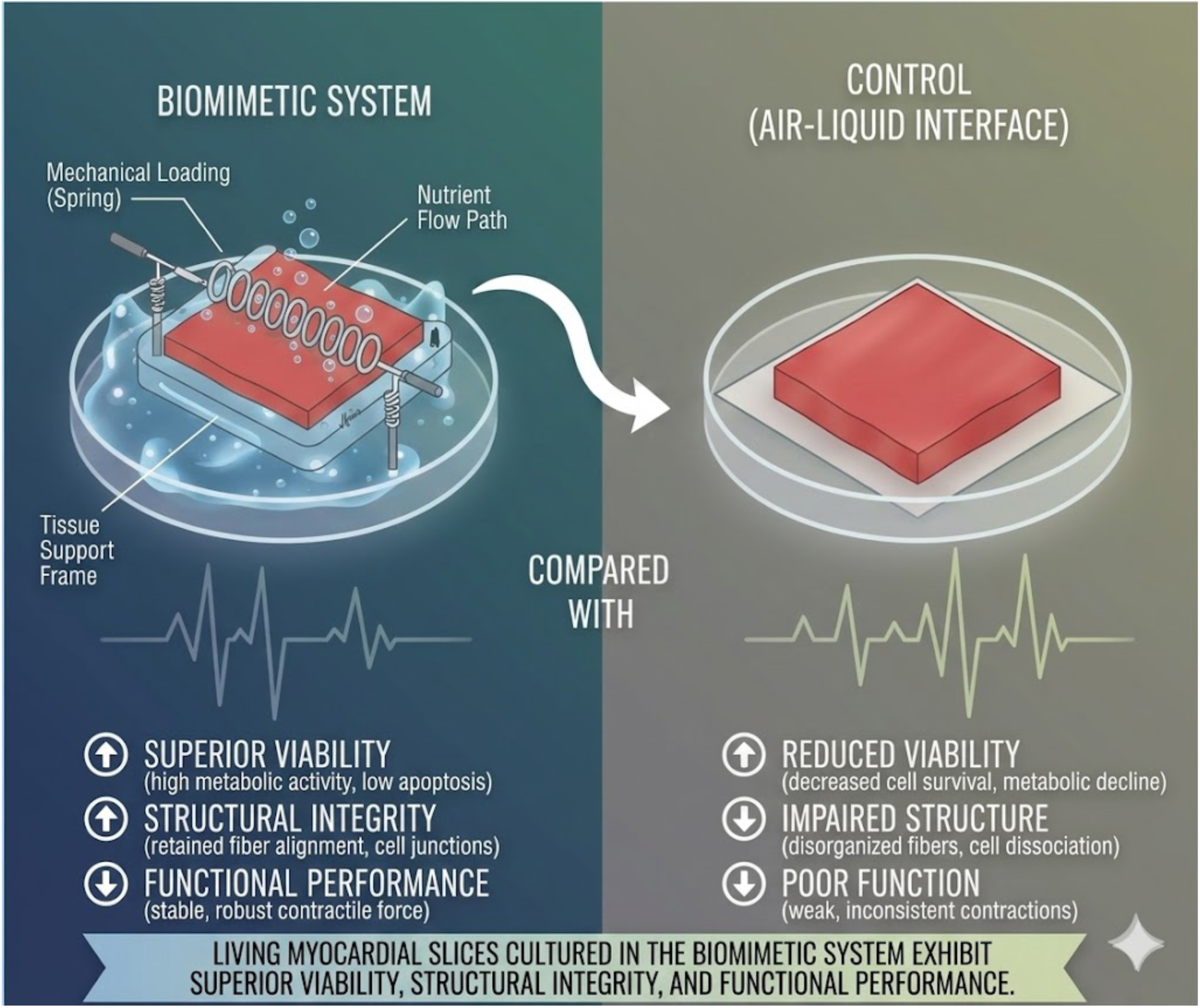

## 1. Introduction

Cardiovascular diseases remain the leading cause of death worldwide, according to the World Health Organization (2025). In 2022, approximately 19.8 million people died from cardiovascular diseases, accounting for an estimated 32% of all global deaths^1–3^. Cardiac fibrosis is a common pathological feature of many cardiovascular conditions, including but not limited to myocardial infarction, heart failure of several aetiologies, genetic- and dilated cardiomyopathies. Furthermore, cardiac fibrosis has also been associated to obesity, diabetes, and aging^4, 5^. Myocardial fibrosis is defined by the accumulation and excessive deposition of extracellular matrix (ECM) proteins within the cardiac interstitium and represents a major component of the response to chronic inflammation.

Fibrotic processes have a complexity that arises in part from the involvement of multiple cell types, including fibroblasts, cardiomyocytes, endothelial cells, and immune cells. Cardiac fibroblasts are widely recognized as key regulators in the cardiac fibrotic response contributing to inflammatory cell recruitment, ECM synthesis and degradation, and scar formation^6^. Fibroblast activation can be triggered by diverse stimuli, including mechanical stress sensed via integrins and mechanosensitive ion channels, neurohormonal factors, inflammatory cytokines and chemokines, and growth factors^5^. Among these, transforming growth factor-β (TGF-β) is one of the most extensively studied pro-fibrotic mediators and is consistently upregulated in experimental models of cardiac fibrosis^4, 5, 7^. TGF-β1 signalling is highly context- and cell-type dependent and promotes fibroblast activation, ECM production, myofibroblast differentiation, increased integrin expression, and a shift toward a matrix-preserving phenotype through the induction of protease inhibitors^4, 5^. The ECM itself actively contributes to fibrosis through embedded signalling molecules and matricellular proteins, including fibronectin, galectins, and proteoglycans such as syndecans^4^. Collagen type I is the most abundant ECM protein in the heart, with collagen type III also present^8, 9^. Increased collagen deposition or crosslinking stiffens the ECM, which is sensed by fibroblasts via mechanosensitive receptors, leading to altered cellular behaviour^4^. Cardiac remodelling is a highly coordinated process involving dynamic interactions between multiple cell types, including cardiomyocytes, fibroblasts, endothelial cells, and immune cells, rather than being driven by fibroblasts alone^4, 5^. Cardiomyocytes can release stress signals that activate fibroblasts, while endothelial cells contribute through endothelial-to-mesenchymal transition and regulation of vascular homeostasis, and immune cells modulate inflammation and profibrotic signalling via cytokine secretion^4, 5, 7^. This complex cellular crosstalk, together with reciprocal interactions with the extracellular matrix, is essential for the initiation and progression of fibrosis, underscoring the need for multicellular models that preserve these interactions^4, 5^.

Despite substantial research efforts, the heterogeneity and complexity of fibrotic remodelling across diverse forms of myocardial injury remain major obstacles to the development of effective therapies ^4^. More refined *in vivo* and *ex vivo* models are needed to identify disease-specific therapeutic targets. One promising approach is the use of living myocardial slices (LMS), which closely recapitulate the native myocardium while requiring only minimal artificial manipulation. LMS are viable, 100–400 µm thick slices of ventricular myocardium generated using a high-precision vibratome^10^. LMS combine advantages of 2-dimentional (D) systems, such as suitability for long-term studies, with those of 3D models, including preserved tissue architecture and ECM, multicellularity, and mature contractile function, while maintaining moderate experimental complexity^11–13^. Compared with whole-heart preparations, papillary muscles, trabeculae, or wedge models, LMS reduce animal use, are easier to manipulate, and allow detailed cellular and subcellular analyses^11–13^. LMS can be prepared from a wide range of mammalian species, including rodents, dogs, pigs, and human cardiac biopsies. Current LMS culture approaches increasingly incorporate mechanical loading and electrical stimulation to enhance physiological relevance, reduce remodelling, and improve tissue viability^11–13^.

Advancing our understanding of cardiac fibrosis and refining experimental models, such as LMS, are critical for the development of effective therapies and improved patient outcomes. In this study, we aimed to establish and evaluate mouse LMS as a novel ex vivo model of myocardial fibrosis.

## 2. Materials and Methods

### Preparation and Culture of LMS

The Experiments were performed on mouse heart tissue of wild type C57BL/6J mice bought from Jackson Laboratory (Charles River, Germany). The animals were kept in the Laboratory Animal Services Center of the University of Zurich in Schlieren under pathogen-free conditions according to the respective license (ZH142/2022). The sample preparation, slicing technique and culturing methods were developed partially by adapting existing protocols ^14–16^. The LMS were cultured *ex vivo* with the biomimetic culture system described by Fischer et al.^17^.

### Extraction and Slicing of the Mouse Heart

Euthanasia was performed using carbon dioxide and according to the respective license (ZH012-2023). After excision the beating heart was rapidly transferred to warm (37°C), heparinized (100 IU) slicing solution (25000 I.E. Heparin-Na, B. Braun, REF: 46613) and gently swirled around to eject the remaining blood from the ventricles. Following this, the heart was transferred to cold (4°C) heparinized solution. The sample was kept cold on ice and quickly transferred to the slicing area. Left ventricular tissue blocks were obtained by dissecting the atria, right ventricle, and the septum. The remaining left ventricular tissue block was glued with the endocardial surface facing upwards onto an agarose block (4%). To slice the heart transversally, the atria were dissected, and the left ventricle was injected with 2% certified Low-Melt Agarose (BioRad, Spain, REF: 1613112) using a syringe (B. Braun, REF: 9166017V) and needle (B. Braun, REF: 4467123). The heart was then embedded in 2% low-melting agarose, with the apex facing upward. The tissue was cut submerged in oxygenated slicing solution at 4 °C and cut by a vibratome (Campden Instruments Ltd, Model 7000smz 2) with a stainless steel (7550/1/SS) or ceramic blade (7550/1/CC), vibrating at 80 Hz with a 2.00 mm amplitude and z-axis error <5 µm. Sections were cut at a thickness of 300 µm, with the blade advancing at 0.03 mm/s (**Scheme 1**).

**Scheme 1.**
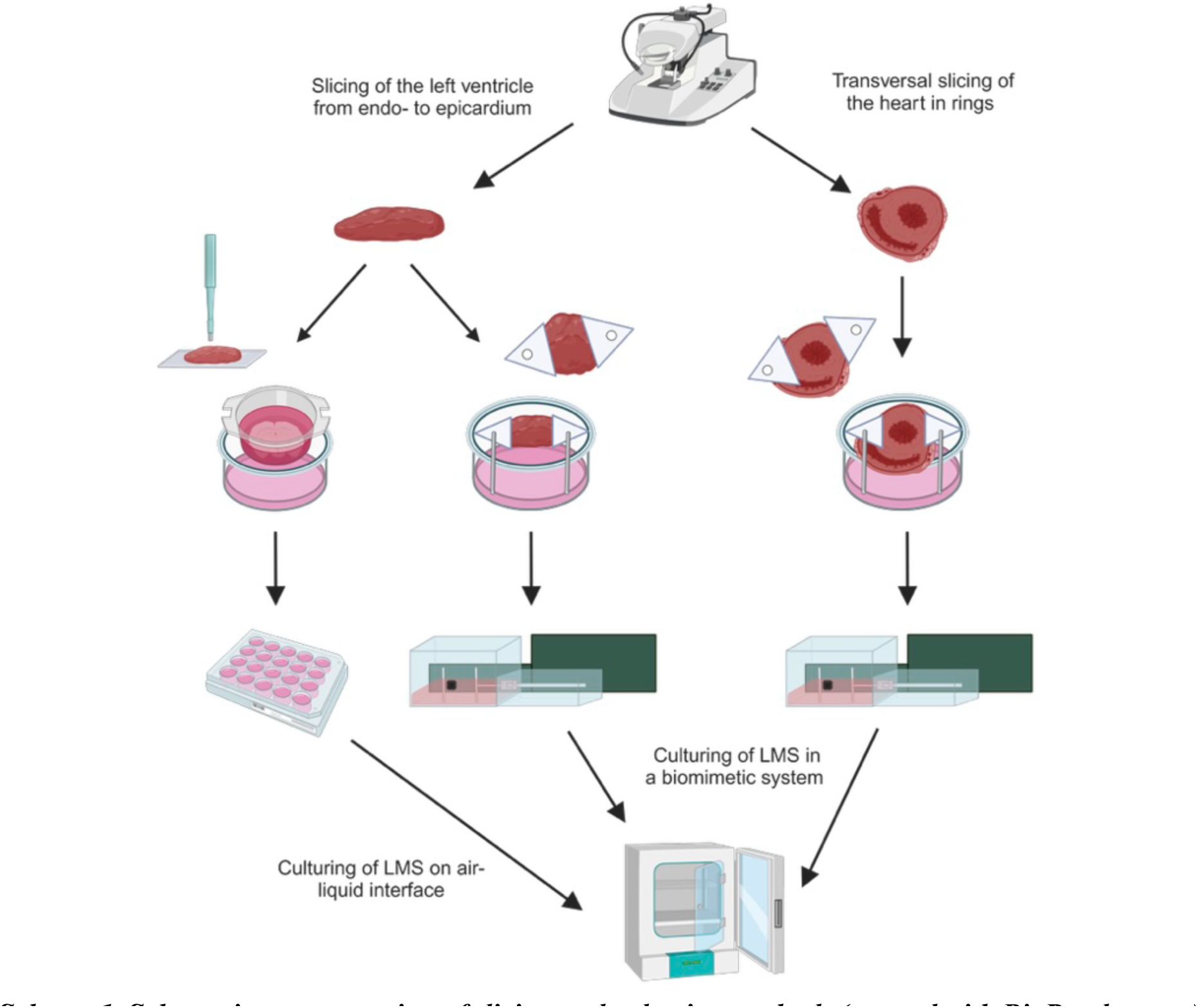
Schematic representation of slicing and culturing methods (created with BioRender.com).

### Culture of the LMS

#### Air-liquid interface system

On day 0, punch biopsies with a diameter of 3 mm were taken from cut section under laminar flow using biopsy puncher (Kai Medical, BPP-30F, Japan) and were placed in culture on inserts (Merck, Ireland, REF: PITP01250 or Thermo Fisher, Denmark, REF: 140627) using the air-liquid interface method. The biopsies were cultured in a standard CO2 incubator at 37C (Binder GmbH, Germany, REF: CB210). After 12-24 h (day 1), medium was exchanged. During the exchange, the Penicillin Streptomycin (PS) concentration was changed from 3% to 1% and the stimulated with 20 ng/mL TGF-β1 (Pepro Tech, USA, REF: 100-21). Subsequently, medium was exchanged every 24h and the relevant tissue samples were stimulated with TGF-β1 (20 ng/mL). On day 5, the tissue culture was terminated, and the tissue was frozen in liquid nitrogen and stored at -80 or used for viability assay (**Scheme 2**).

**Scheme 2.**
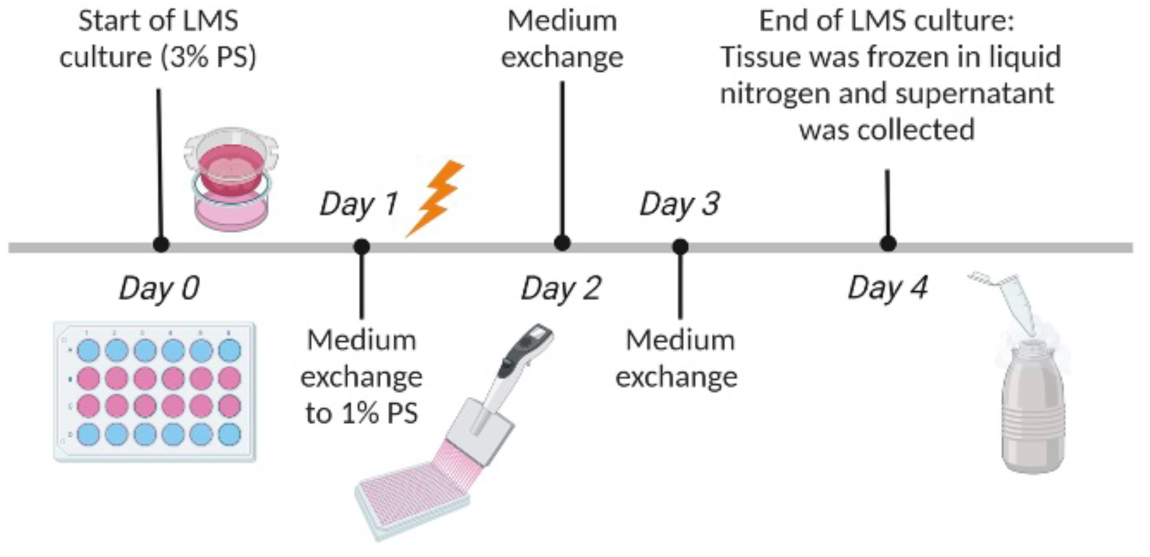
Experimental timeline for air-liquid interface culture of LMS (created with BioRender.com).

#### Biomimetic MyoDish system

The LMS derived from isolated left ventricular tissue blocks, both axially and ring-shaped transversely cut, were cultured in the MyoDish 1 Tissue Culture System (InVitroSys GmbH, Model MD-1.2). inside a standard incubator (Binder GmbH, CO2 Incubator, REF: CB210) plastic triangles were dipped in Histoacryl® glue (Kaladent, REF: 2560107) and affixed to the tissue. The sample was placed in a culture chamber, and the preload was set to either 300 or 1200 μN. Starting on day 0 half of the tissue samples were cultivated in medium supplemented with 20 ng/mL TGF-β1. Subsequently, the medium was changed every 48 hours and the relevant tissue samples were stimulated with TGF-β1 (20 ng/mL). The tissue was paced at 50mA and 60bpm. Measurements of force-frequency relationship, stimulation threshold, post-rest potential, and refractory period were taken every 24 hours. On day 6, the tissue culture was terminated, and the tissue was frozen in liquid nitrogen and stored at -80° (**Scheme 3**).

**Scheme 3.**
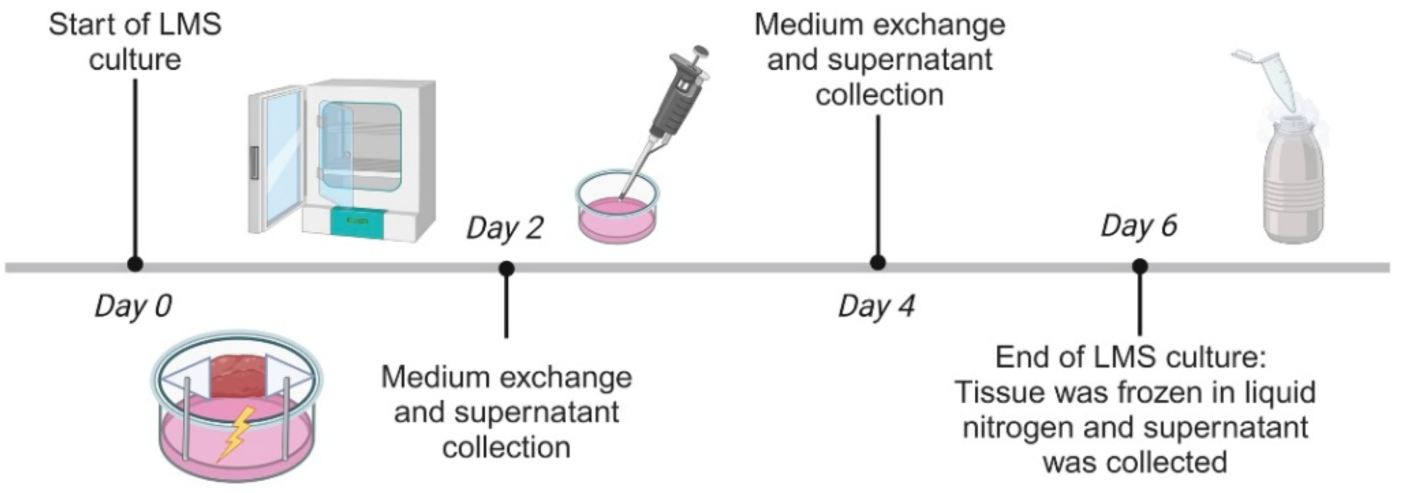
Experimental timeline for biomimetic MyoDish system of (created with BioRender.com).

#### 1.1.1 Viability assay using LIVE/DEAD™ Viability/Cytotoxicity Kit for mammalian cells

The LIVE/DEAD™ Viability/Cytotoxicity Kit for mammalian cells (Thermo Fisher Scientific, REF: L3224) was used to stain the myocardial tissue samples and assess their viability. Samples were incubated for 30 minutes. Images were acquired using the Leica SP8 inverse confocal laser-scanning microscope (Center for Microscopy and Image Analysis at the University of Zurich). During the transport and imaging, the tissue remained inside a sufficient amount of PBS to prevent the tissue from drying out. The dead control was established by a 10 minute incubation in 80% ethanol solution. The signal intensities were quantified using the ImageJ software. Staining intensity is presented in arbitrary units (AU) representing the ratio of the signal from the death to the alive tissue.

#### 1.1.2 MTT viability assay

Viability measurement was performed using Thiazolyl Blue Tetrazolium Bromide (MTT) (Merck REF: M5655-100MG) dissolved in slicing solution at a concentration of 0.5 mg/mL. In a 48-well plate, one sample per well was treated with 200 µL of the MTT solution for an incubation period of 25 minutes at 37°C. Subsequently, the samples were washed with slicing solution for 5 minutes at room temperature. Following this, the samples were incubated in DMSO (Carl Roth GmbH, REF: A994.2) for 30 minutes at 37°C, with an additional stirring step after 20 minutes. Post-incubation, absorbance readings of the DMSO solution were performed at 555 nm using the Synergy HT microplate reader using the Gen5 software (BioTek, version 2.04). The tissue was air-dried and weighed.

#### 1.1.3 Lactate dehydrogenase (LDH) Cytotoxicity Assay

LDH levels were determined using the LDH-Glo™ Cytotoxicity Assay (Promega, REF: J2380) on samples diluted 1:50 in storage buffer (200mM Tris-HCl, 10% Glycerol, 1% BSA) according to the manufacturer’s instructions. The signal was acquired using the Synergy HT microplate reader using the Gen5 software (BioTek, version 2.04). The tissue was air-dried and weighed.

### 1.2 Quantitative RT-PCR

For one replicate, four to eight punch biopsies or two slices were pooled. The tissue was homogenized with Bertin Technologies Minilys homogenizer at 5000 rpm for a duration of 5-10 seconds, and total RNA was isolated using the RNeasy Mini Kit (Qiagen, REF: 74106) or TRI Reagent®. The RNA content was measured using a NanoDrop 2000 Spectrophotometer (Thermo Fisher Scientific). cDNA synthesis was performed using the BioRad T100 Thermal Cycler. The cDNA was amplified using the Agilent Technologies Stratagene Mx3005P. Gene expression was detected using the SYBR green GoTaq qPCR Master mix (Promega, REF: A6002 Dubendorf, Switzerland) mixed with Nuclease free water (Promega, REF: P119E) and oligonucleotides complementary to transcripts of the analysed genes (**Table 1**). Transcript levels of mouse RPLP0 and mouse GAPDH were used as endogenous reference, and relative gene expression was analysed using the 2-ΔΔCT method ^18^. Mice forward and reverse primer pairs used for RT-qPCR were acquired from Microsynth (**Table 2**).

**Table 1.** Components used to generate cDNA.

| Name of component | Manufacturer and Reference Number | Country |
| --- | --- | --- |
| dNTPs 10 mM | Thermo Fisher Scientific, REF: 362275 | UK |
| 10x PCR Buffer II | Thermo Fisher Scientific, REF: 4486220 | Lithuania |
| RNase free H2O | Invitrogen, REF: 10977-049 | UK |
| Magnesium Chloride (50mM) | Sigama-Aldrich, REF: M1028-10XML | USA |
| MultiScribe Reverse Transcriptase | Thermo Fisher Scientific, 50 U/μL, REF: 4308228 | USA |
| Random Hexamers (50μM) | Thermo Fisher Scientific, REF: 100026484 | USA |
| RNase Inhibitor | Thermo Fisher Scientific, 20 U/μL, REF: N8080119 | Lithuania |

**Table 2.**
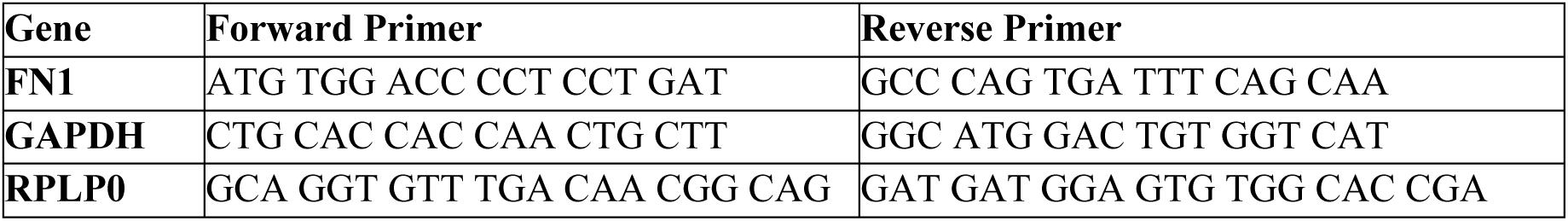
Mice forward and revers primer pairs used for RT-qPCR.

### 1.3 ELISA

Supernatants from individual tissue slices were collected during each medium change and stored at −80°C. Pro-collagen Iα1 levels were measured using the Mouse Pro-Collagen I Alpha 1 Matched Antibody Pair Kit (Abcam, REF: ab216791) following the manufacturer’s instructions. Optical density was measured at 450 nm using a Bio-Tek Synergy HT microplate reader (Winooski, Vermont, USA) and Gen5 software (BioTek, version 2.04). Pro-collagen Iα1 levels were determined from a standard curve.

### 1.4 Histology and Immunohistochemistry

Myocardial tissue was washed in PBS and fixed overnight in 4% paraformaldehyde at 4°C, followed by transfer to 50% ethanol for at least 12 hours at 4°C. Tissues were dehydrated using a graded ethanol series (80%, 96%, 100%), incubating for 1 hour at each concentration, and then cleared in xylene with two 1-hour incubations. Tissue was embedded in paraffin at 56°C, cooled at -20°C, and sectioned into 2–3 µm slices using a microtome. Sections were mounted on Superfrost Plus microscope slides (Thermo Scientific) and dried overnight at 58°C. Sirius Red staining (Direct Red 80, Sigma-Aldrich, REF: 365548) was performed for 3 minutes to visualize collagen I and III fibers, with nuclei counterstained using a 1:1 ratio of Weigert A and B reagents (Artechemis, REF: US-081 and US-082) for 8 minutes. Hematoxylin and eosin (H&E) staining was performed by incubating in Mayer’s hematoxylin (Artechemis, REF: US-020) for 10 minutes, followed by eosin (BioSystems, REF: 84-0012-00) for 3 minutes. Deparaffinization was done using two washes in xylene for 2 minutes, followed by immersion in a descending alcohol series (100%, 100%, 96%, 80%). After staining, the sections were dehydrated in a reverse order of alcohols.

For immunohistochemistry, the Leica BOND-MAX™ automated system (Leica Biosystems) was used with the BOND Polymer Refine Detection System with DAB chromogen (Leica, REF: DS9800) Antigen retrieval was performed using Tris-EDTA (pH 9) or citrate buffer (pH 6), as specified for each antibody (Table 3). Endogenous peroxidase activity was blocked using hydrogen peroxide, followed by incubation with primary and secondary antibodies (Table 3). Nuclei were counterstained with hematoxylin. Images were captured using an Olympus BX53 microscope with a DP80 camera and cellSens standard imaging software (version 1.17) or the Zeiss Axio Scan.Z1 slide scanner, supported by the Center for Microscopy and Image Analysis, University of Zurich.

**Table 3.** Primary and secondary antibodies used for immunohistological staining on mouse LMS.

| Antibody | Dilution | Clone | Antigen Retrieval | Manufacturer & Reference Number |
| --- | --- | --- | --- | --- |
| **Anti-α-SMA** | 1:2000 | E184 | Tris EDTA for 30 min at 100°C | Abcam, REF: 132575 |
| **Anti-Vimentin** | 1:10000 | EPR3776 | Tris EDTA for 30 min at 100°C | Abcam, REF: ab92547 |
| **Anti-von Willebrand factor** | 1:100 | EPR25069-131 | Tris EDTA for 30 min at 100°C | Abcam, REF: ab287962 |
| **Anti-Periostin** | 1:2000 | Polyclonal | Tris EDTA for 30 min at 100°C | Abcam, REF: 14041 |
| **Anti-Troponin-T** | 1:10000 | 13-11 | Citrate buffer for 20 min at 95° | Thermo Fisher Scientific (Invitrogen), REF: MA5-12960 |
| **Anti-gp38** | 1:10000 | eBio81.1(8.1.1) | Citrate buffer for 20 min at 95° | Thermo Fisher Scientific; REF: 14-538182 |
| **Rabbit anti-Hamster IgG (H+L)** | 1:1000 + 2% mouse serum |  |  | Thermo Fisher Scientific, REF: A18891 |

**Table 4.** Solutions used.

| **Solution** | **Components** | **Reference** | **Country** |
| --- | --- | --- | --- |
| **Slicing solution / Tyrode's solution** | 30mM 2,3-Butanedione monoxime (BDM) | Sigma-Aldrich, REF: B0753-25G | India |
|  | 140mM Sodium chloride | Carl Roth GmbH & Co. KG, REF: 3957.2 | Germany |
|  | 6mM potassium chloride | Merck, REF: 1.04936.1000 | Germany |
|  | 10mM glucose | Sigma, REF: G-5400 | USA |
|  | 10mM HEPES | Carl Roth GmbH & Co. KG, REF: 6763.2 | Germany |
|  | 1mM magnesium chloride solution (1M) | Merck, REF: M1028-100ML | USA |
|  | 1.8mM calcium chloride solution (1M) | Fluka, REF: 21115 | Switzerland |
| | In dH<sub>2</sub>O at pH 7.4 filtered using 0.22 μm pore filter | TPP Techno Plastic Products AG, REF: 99255 | Switzerland |
| **Medium** | Medium-199 | Sigma-Aldrich, M2154 | UK |
| | ITS Liquid Media Supplement (100×) | Sigma-Aldrich, REF: I3146 | USA |
|  | Penicillin Streptomycin | Gibco by Life Technologies, REF: 15070-063 | India |
|  | Fungizone® Amphotericin B | Gibco by Life Technologies, REF: 15290-026 | Israel |
| | Phenylephrine-hydrochloride 50μM | Sigma-Aldrich, REF: P6126 | China |
|  | L-ascorbic acid 2-phosphate aequimagnesium salt hydrate 0.18 mM | Sigma-Aldrich, REF: A8960-5G | Japan |
|  | 0.1 % 2-Mercaptoethanol (50mM) | Gibco by Life Technologies, REF: 31350-010 | Scotland |

### 1.5 Statistical Analysis

Statistical tests and analysis were performed using GraphPad Prism software (version 10). P values were considered significant when < 0.05. Fold changes were calculated relative to the appropriate control. In general, the fold changes were calculated to the average of the control group per individual experiment. However, in the experiments using the pacing system, the control was the tissue sample itself on day 0 or 1 for a first normalization. When comparing the TGF-β1 stimulated group to the control group the fold changes were calculated additionally to the average of the relevant control group.

### 1.6 Materials

## 2 Results

### 2.1 Comparison of LMS viability in air-liquid interfaces and biomimetic system (MyoDish)

To assess LMS viability, we used the LIVE/DEAD™ Viability/Cytotoxicity Kit, which distinguishes live from dead cells, with dead cells labeled by ethidium homodimer-1 (**Figure 1A)**. The application of this assay for evaluating viability in myocardial tissue slices has been previously established, and dead cells are measured by the ratio of red:green signal^13^. Quantitative analysis revealed pronounced differences between the dead control and viable samples, where the viable samples show a significant reduction in the red:green signal (**Figure 1B**). Moreover, the ratio of red to green fluorescence increased progressively over the culture timeline, indicating a gradual decline in tissue viability (**Figure 1C**). Due to a reduction of viability at day five, it was concluded that the murine LMS can be maintained on the air-liquid interface for a maximum of four days. However, it should be noted that the assay is limited by incomplete dye penetration across the full tissue thickness. Notably, LMS cultured in the biomimetic MyoDish system could not be assessed for viability using the LIVE/DEAD™ Viability/Cytotoxicity Kit, as tissue folding after culture in this system prevented reliable staining and analysis.

**Figure 1.**
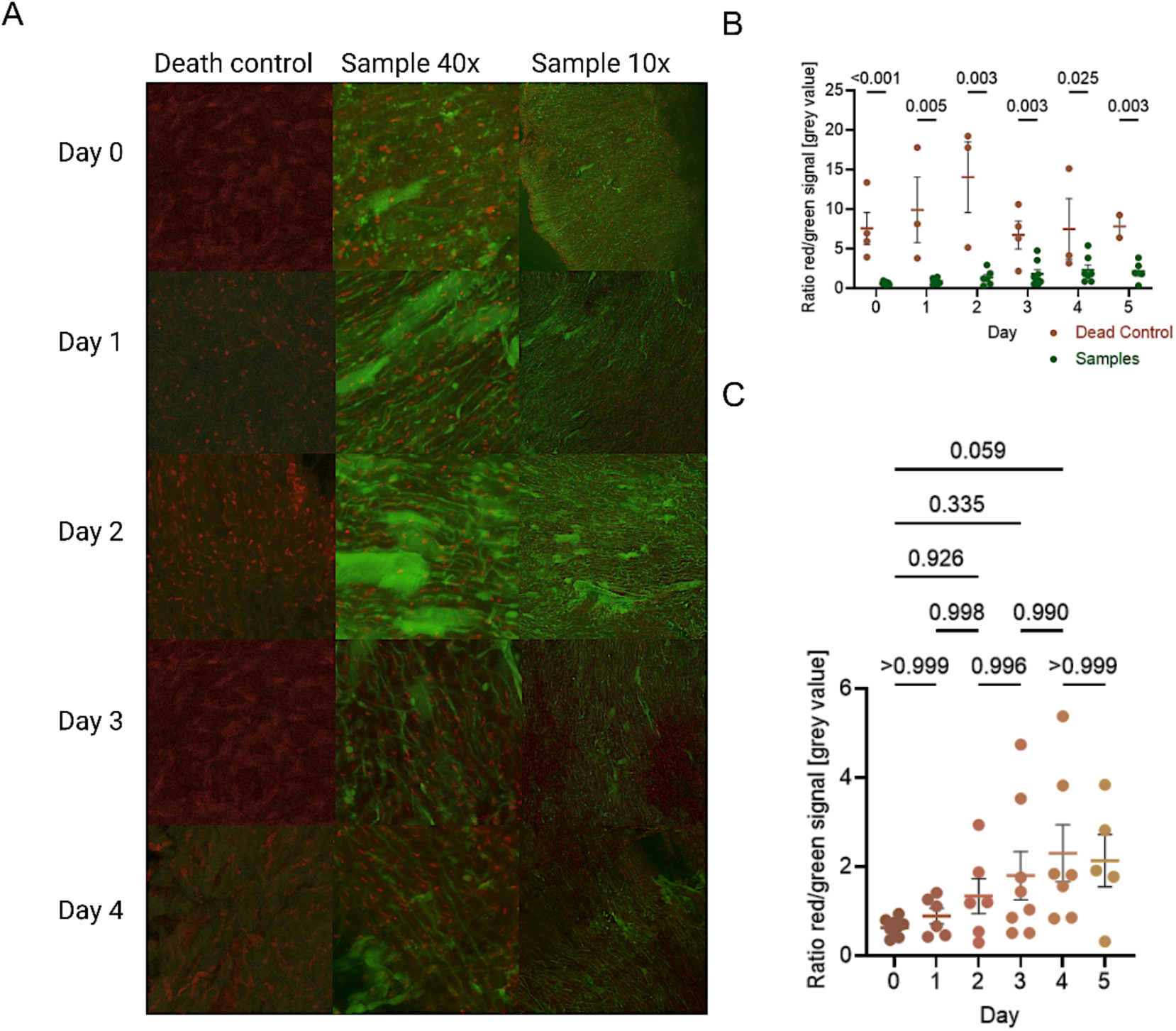
Viability determination of mouse LMS using LIVE/DEAD™ Viability/Cytotoxicity Assay. (**A**) Representative images of viability staining on LMS from air–liquid interfaces system of day 0-4 in culture using LIVE/DEAD™ Cell Imaging Kit. Stained samples are shown at a 10-fold and 40-fold magnification. The dead control (DC) is displayed at 40-fold magnification. (Red: marking the dead or dying cells; green: staining only the cells with active metabolism). (**B**) Quantification of the fluorescent signal red divided by green of the gray value from the summed-up picture of the z-stack. Comparing the ratio of the dead control (Control, red) to the sample (Sample, green) using a multiple, unpaired, two-tailed parametric t-test employing the two-stage setup (Benjamini, Krieger, and Yekutieli) with a false discovery rate of 1%, hence showing the q-values. Data from four different experiments; n = 2-9. (**C**) Signal ratio of red to green signal of the gray value from the summed-up picture of the z-stack of only the samples. An ordinary One-way ANOVA using Šídák’s multiple comparisons test was applied. Data from four different experiments; n = 2-9.

To enable a direct comparison of myocardial tissue viability between air-liquid interface culture and the biomimetic MyoDish system, viability of mouse LMS was assessed using the MTT assay. Measurements were performed on day 2 and again on day 6 at the end of the culture period. A time-dependent decrease in viability was observed in both TGF-β1-stimulated and unstimulated LMS cultured at the air-liquid interface as well as in the biomimetic MyoDish system (**Figure 2A-B**). Notably, LMS maintained in the biomimetic MyoDish system exhibited higher viability in control conditions at day 2, which was reduced following TGF-β1 stimulation and culture time (**Figure 2B**). In contrast, LMS cultured on the air-liquid interface displayed relatively low viability already at day 2 (**Figure 2A**). LMS viability was further evaluated using the LDH-Glo™ Cytotoxicity Assay. For this analysis, LMS were electrically paced in the biomimetic MyoDish system, with each stimulation frequency applied for at least 10 seconds under a preload of either 300 µN (top) or 1200 µN (bottom), targeting approximately 40 beats per tested frequency. Most cell death occurred after four days of culture, with neither increased preload nor TGF-β1 stimulation altering LDH release compared with control conditions (**Figure 2C**).

**Figure 2.**
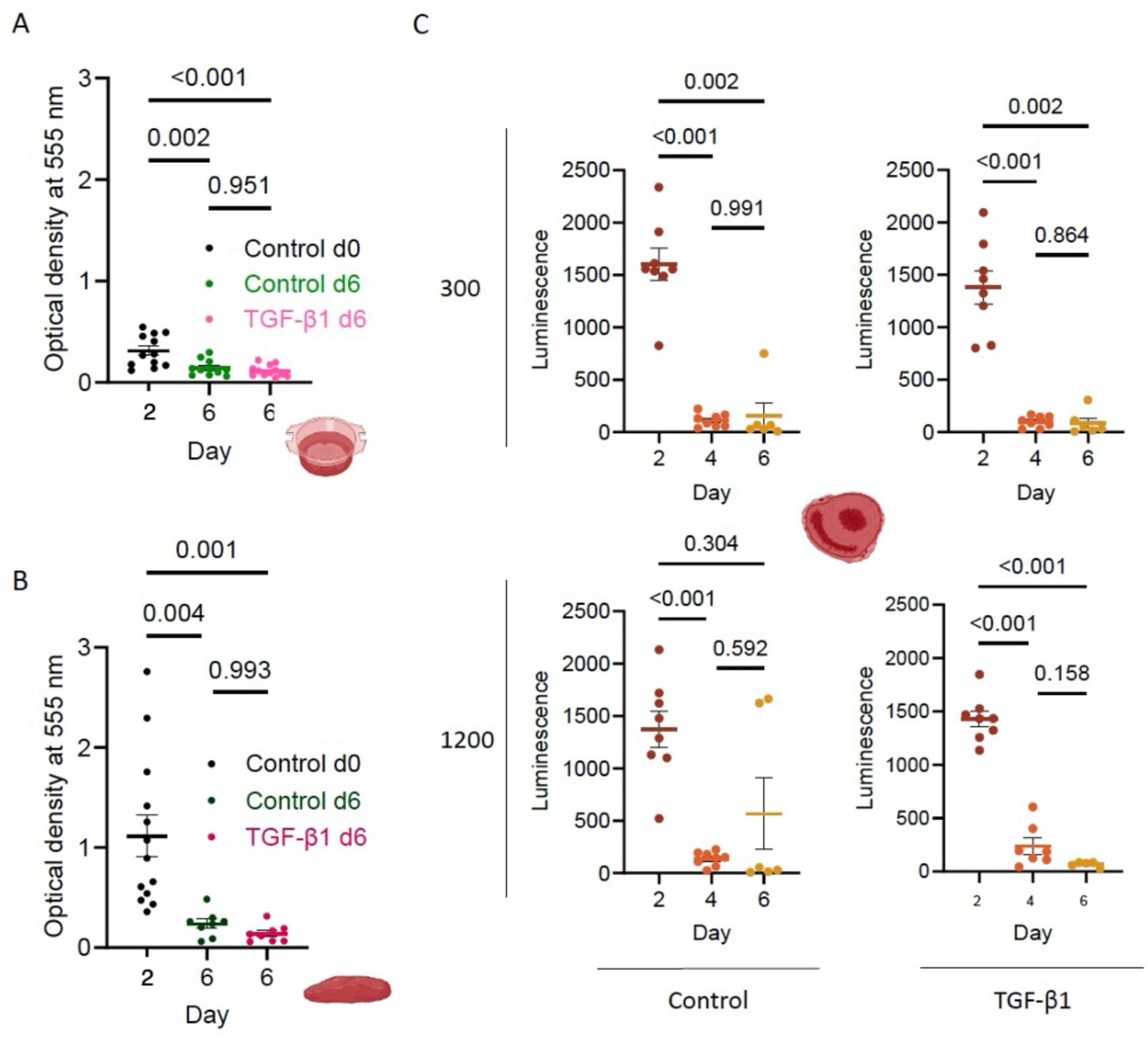
Viability of LMS decreases with culture duration across all conditions. (**A-B**) A progressive decline in LMS viability over time was detected using the Thiazolyl Blue Tetrazolium Bromide (MTT) assay. LMS cultured at the air-liquid interface (**A**) and LMS prepared from the same hearts using an identical slicing protocol but maintained in the biomimetic MyoDish system (**B**), both showed a time-dependent reduction in metabolic activity. Statistical analysis was performed using ordinary one-way ANOVA with Šídák’s multiple comparisons test (n = 8-13). (**C**) Cytotoxicity assessment using the LDH-Glo™ Cytotoxicity Assay showed that most cell death occurred within the first four days of culture. Data shown are from transversely cut, ring-shaped LMS cultured in the biomimetic MyoDish system under a preload of either 300 µN (top) or 1200 µN (bottom). Mixed effect analysis, Šídák’s multiple comparisons test, n=4-7.

### 2.2. Assessment of contractile function in LMS cultured in the biomimetic MyoDish system

Using the biomimetic MyoDish culture system, contractile force is measured as the lateral displacement of a spring wire during LMS contraction, enabling continuous and detailed functional monitoring. The system also supports pre-programmed stimulation protocols, allowing regular assessment of parameters such as stimulation threshold, force-frequency relationships, and post-rest potentiation. Contractile function of LMS cultured in the biomimetic MyoDish system was assessed by measuring contraction amplitude, which is recorded by the sensor and directly reflects contractile force. To minimize background noise during amplitude measurements, the rocking board was switched off. Measurements were performed every 24 h. As expected, a high degree of variability in contraction amplitude was observed between individual tissue slices, likely reflecting differences in slice size and their anatomical origin within the heart. Following an initial adaptation period of approximately 24 h, contraction amplitudes stabilized across cultures (**Figure 3**).

**Figure 3.**
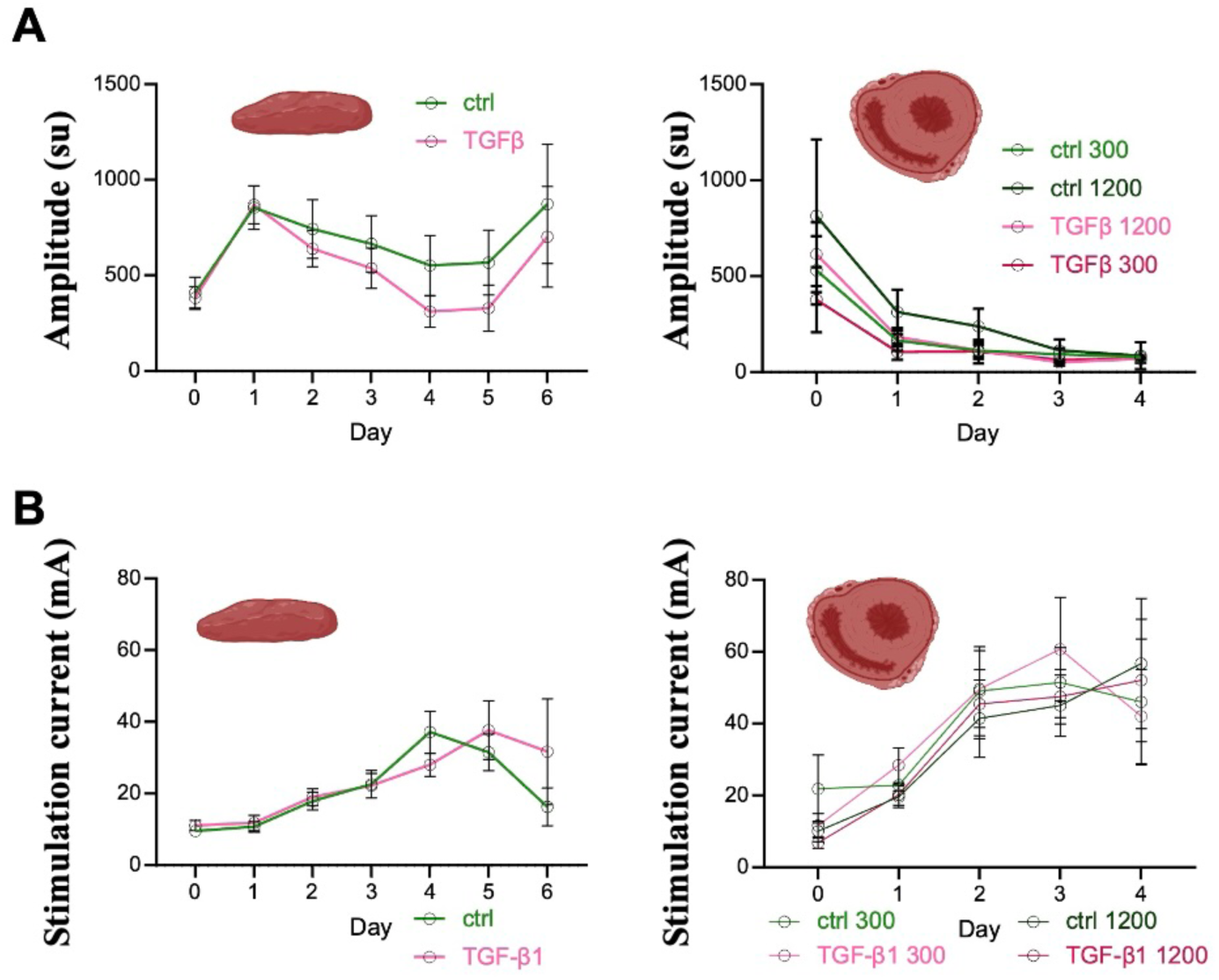
LMS cultures prepared from isolated left ventricular tissue blocks exhibit superior quality compared with LMS obtained by transverse slicing of whole embedded hearts. (**A**) LMS derived from isolated left ventricular tissue blocks display more stable contraction force throughout the culture period compared to LMS prepared by transverse slicing of embedded hearts. Contraction amplitudes, which directly reflect contractile force, were measured on each viable day of culture. No significant differences between the two groups using an unpaired, multiple t-test with a FDR desired (Q) 1% using the Two-stage step-up method by Benjamini, Krieger, and Yekutieli. Pulled data from four experiments, meaning up to 16 samples per group. Abbreviations: sensor unit (su). (**B**) LMS prepared by transverse slicing of embedded hearts show an earlier increase in stimulation threshold during culture compared to LMS derived from isolated left ventricular tissue blocks (n=6-10, n=3-10). Stimulation threshold measured on each day possible in culture. No significant differences between the two groups using an unpaired, multiple t-test with a FDR desired (Q) 1% using the Two-stage step-up method by Benjamini, Krieger, and Yekutieli. Pulled data from four experiments, meaning up to 16 samples per group. Statistical analysis was performed by GraphPad Prism (version 10). Abbreviations: ctrl = control unstimulated, TGF-β1 = TGF-β1 stimulated (20 mg/mL).

Functional testing was performed under electrical stimulation, with each stimulation frequency applied for a minimum of 10 s and a maximum of 300 or 1200 s, targeting approximately 40 beats per frequency. TGF-β1 was applied as a pro-fibrotic stimulus. LMS cultures prepared from isolated left ventricular tissue blocks showed superior functional performance compared with those obtained by transverse slicing of whole embedded hearts. Specifically, LMS derived from isolated left ventricular tissue blocks maintained a more stable contraction force throughout the culture period compared to LMS generated by transverse slicing of embedded hearts (**Figure 3A**). However, prolonged stimulation at higher frequencies (up to 1200 s) in combination with TGF-β1 treatment resulted in a progressive reduction in contraction amplitude over time in both LMS culture variants (**Figure 3A**). In contrast, LMS generated by transverse slicing of embedded hearts exhibited an earlier increase in stimulation threshold during culture compared to LMS derived from isolated left ventricular tissue blocks, indicating a more rapid decline in functional responsiveness (**Figure 3B**). TGF-β1 treatment and prolonged stimulation at higher frequencies (up to 1200 s) further contributed to a modest increase in stimulation threshold at later time points (days 5-6; **Figure 3B**). Overall, the pronounced variability in contraction amplitude among LMS likely reflects inherent biological heterogeneity of myocardial tissue, emphasizing the importance of standardized tissue preparation for functional studies.

In the context of heart failure, contractile dysfunction is often associated with impaired excitation– contraction coupling and altered calcium (Ca²⁺) homeostasis. Post-rest potentiation (PRP) is linked to Ca²⁺ accumulation within the sarcoplasmic reticulum during pauses in electrical stimulation and reflects the tissue’s capacity to store and release calcium. Accordingly, PRP measurements provide insight into myocardial functional integrity and overall tissue health. To assess post-rest potentiation (PRP), we applied a predefined stimulation protocol consisting of progressively increasing pause durations, followed by 30 s of electrical stimulation at 50 mA and 60 bpm to allow functional recovery. Upon resumption of stimulation, the amplitudes of the first one to three contractions were recorded.

**Figure 4** illustrates the relationship between stimulation pause duration and contraction amplitude from day 0 to day 4 of LMS culture. Data are presented as fold changes in amplitude following stimulation pauses of 1-120 s, normalized to contractions measured after a 2 s pause at a baseline frequency of 60 bpm, for LMS derived from isolated left ventricular tissue blocks (**Figure 4A**) and LMS generated by transverse slicing of embedded hearts (**Figure 4B**). On day 0, contraction amplitude increased after longer pauses (12 and 120 s) compared with baseline stimulation; however, this effect diminished over the course of culture (**Figure 4A-B**). No differences were observed between TGF-β1-stimulated and control groups at either day 0 or day 4 across all experimental conditions (**Figure 4A-B**).

**Figure 4.**
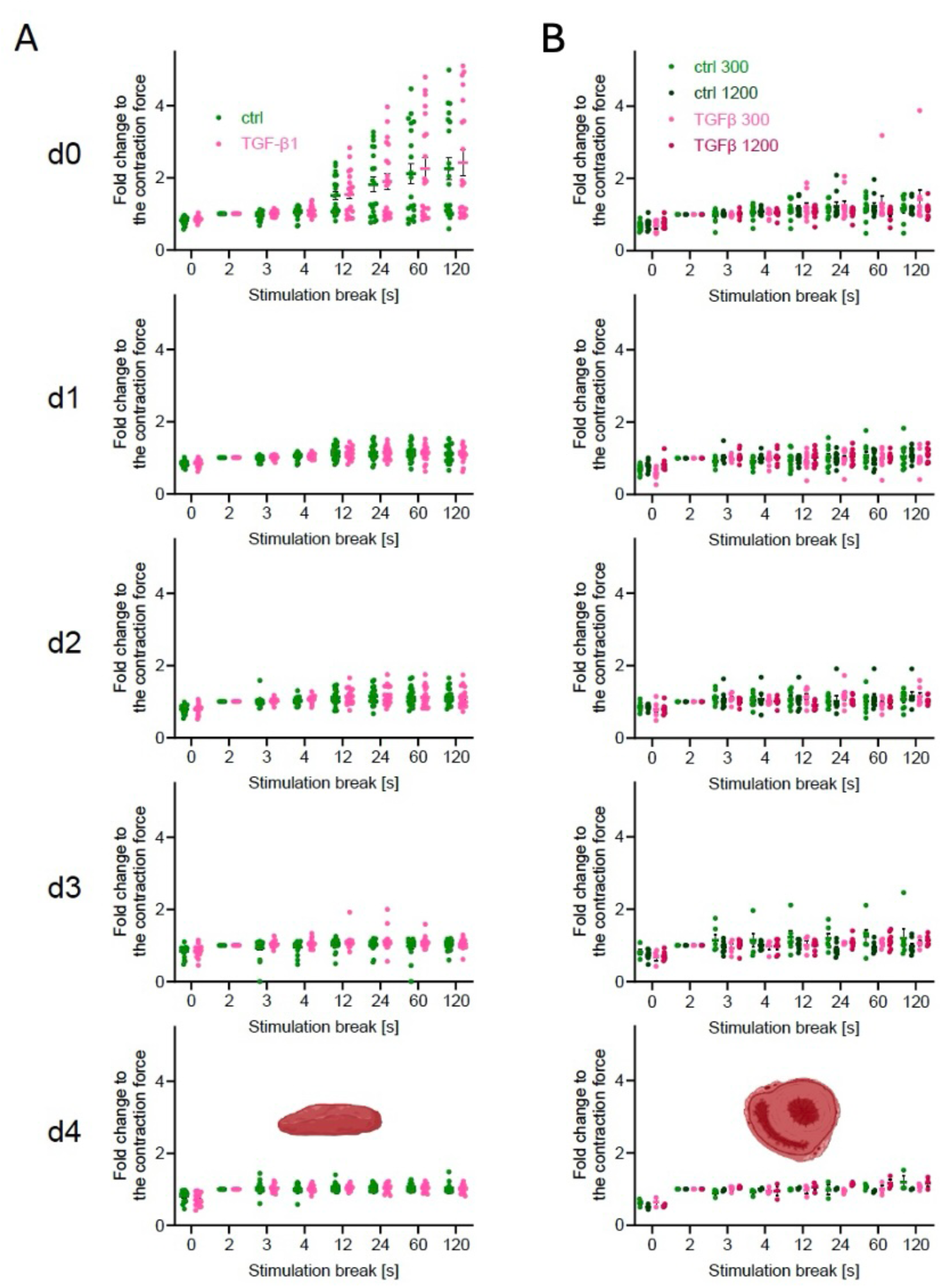
Interruption of electrical stimulation induces contractile potentiation in LMS. (**A**) Fold change in contraction amplitude following stimulation pauses of varying duration (1-120 s), normalized to amplitudes measured without a stimulation break. Data were pooled from four independent experiments, comprising 9-16 LMS derived from isolated left ventricular tissue blocks. TGF-β1 was applied as a profibrotic stimulus. (**B**) Fold change in contraction amplitude following stimulation pauses of varying duration (1-120 s), normalized to amplitudes measured without a stimulation break. Data were pooled from four independent experiments, comprising 9-16 LMS generated by transverse slicing of embedded hearts. Functional testing was performed under electrical stimulation, with each stimulation frequency applied for a minimum of 10s under a preload of either 300 µN (top) or 1200 µN (bottom), targeting approximately 40 beats per frequency. TGF-β1 was applied as a profibrotic stimulus.

Contraction amplitude was increased with longer stimulation pauses across both unstimulated and TGF-B1-stimualted conditions. A significant effect was observed between the culture days (day 0 vs day 4) for only LMS derived from isolated left ventricular tissue blocks (**Figure 5A**). In contrast, for LMS obtained by transverse slicing of embedded hearts, no significant differences between day 0 and day 3 were detected (**Figure 5B**). Together, across both preparation methods, PRP increased with longer stimulation breaks but progressively declined with time in culture, indicating a gradual loss of functional calcium handling capacity.

**Figure 5.**
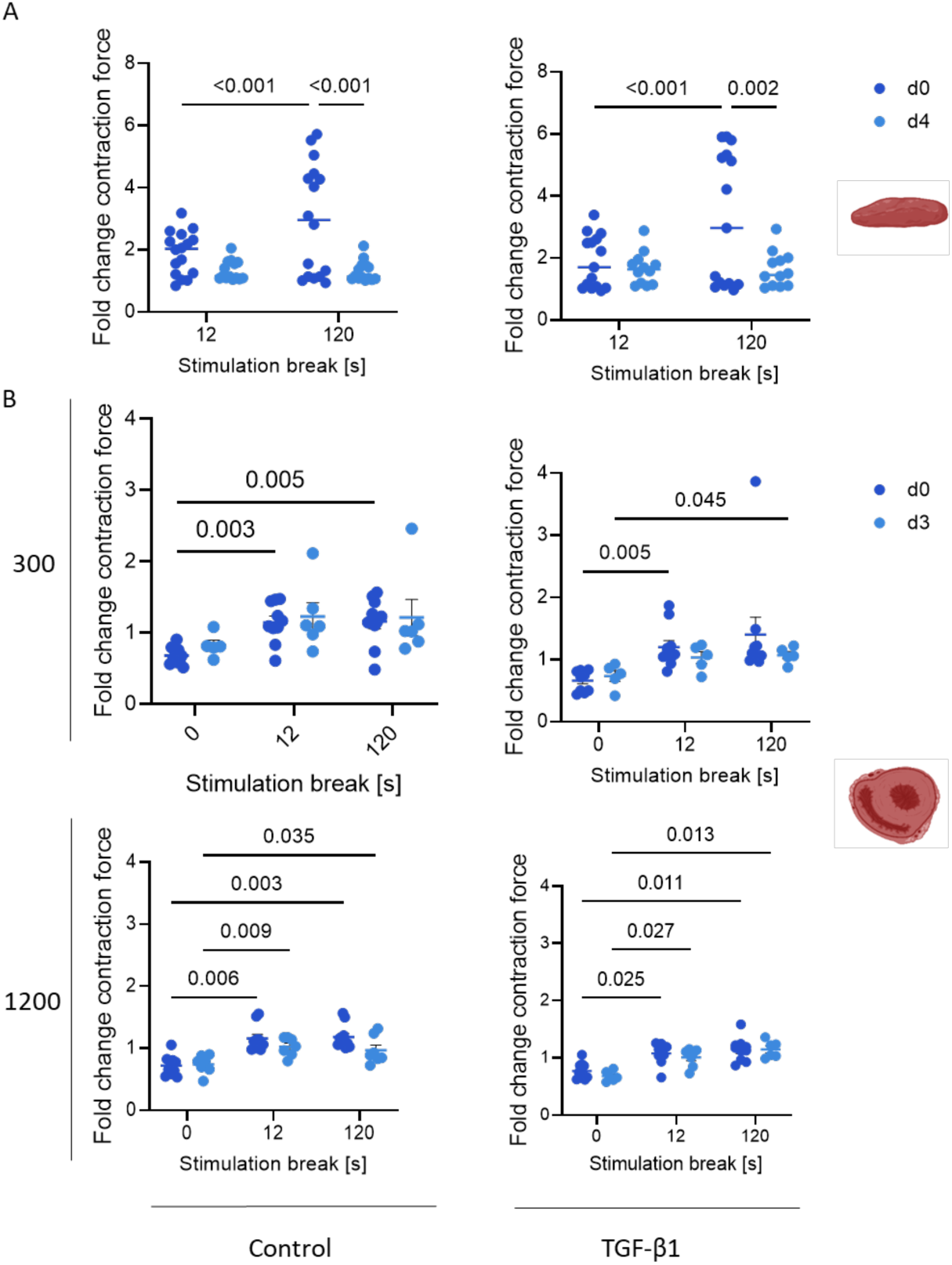
Post-rest potentiation relationship of LMS changes with increasing pacing frequency. (**A**) Fold change in contraction force following stimulation pauses of varying duration (1–120 s), normalized to amplitudes measured without a stimulation break. Data were pooled from four independent experiments, comprising 9-16 LMS derived from isolated left ventricular tissue blocks. TGF-β1 was applied as a profibrotic stimulus. Two-way ANOVA Fishers LSD test comparing the means of the days to each other as well as the stimulation breaks displaying mean, SEM interleaved scatter. (**B**) Fold change in contraction force following stimulation pauses of varying duration (1–120 s), normalized to amplitudes measured without a stimulation break. Lines are drawn as visual guides only. Data were pooled from four independent experiments, comprising 9-16 LMS generated by transverse slicing of embedded hearts. Functional testing was performed under electrical stimulation, with each stimulation frequency applied for a minimum of 10s under a preload of either 300 µN (top) or 1200 µN (bottom), targeting approximately 40 beats per frequency. TGF-β1 was applied as a profibrotic stimulus. Two-way ANOVA Tukey’s (ii & iii), Mixed-effects model (REML) analysis (i & iv), Tukey’s multiple comparisons test, testing the means of all simulation breaks against each other and the means of both conditions at individual stimulation stops against each other.

The force-frequency relationship (FFR) of LMS changed over the course of tissue cultivation (**Figure 6**). A negative FFR has been associated with altered intracellular Ca²⁺ handling and reduced T-tubular density^19^, therefore, assessment of the FFR provides an indirect measure of calcium-handling integrity and overall tissue functionality. To evaluate the relationship between electrical stimulation frequency and contractile force, a pre-defined stimulation protocol was applied in which pacing frequency was gradually increased from 12 to 210 bpm. Each frequency was maintained for a minimum of 10 seconds and a maximum of 300 seconds, targeting approximately 40 contractions per frequency. Contractile force was quantified as contraction amplitude. **Figure 6** depicts the FFR measured from day 0 to day 4 of culture in LMS from isolated left ventricular tissue blocks (**Figure 6A**) or LMS prepared either by transverse slicing of embedded hearts (**Figure 6B**). No differences were observed between TGF-β1-stimulated and control groups across all experimental conditions (**Figure 6A-B**).

**Figure 6.**
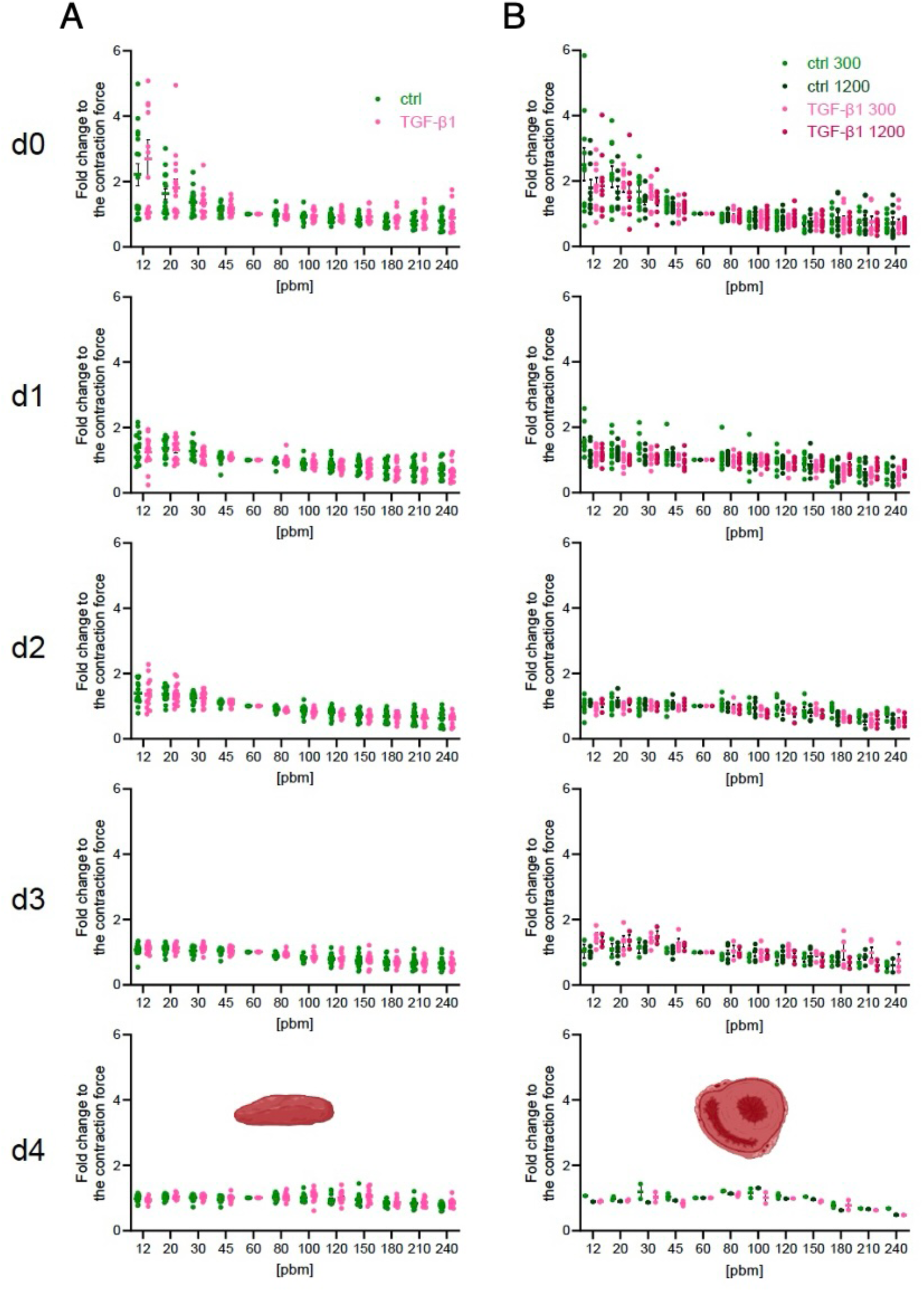
Force-frequency relationship of LMS changes with increasing time in culture. (**A**) Fold change of the amplitude shown at different frequencies of stimulation 12-240 bpm to the amplitude measured at 60 bpm (each sample to itself) for. Pulled data from 4 experiments together leading to data from 6-16 LMS derived from isolated left ventricular tissue blocks. TGF-β1 was applied as a profibrotic stimulus. (**B**) Fold changes calculated of individual sample to itself on amplitude measurements at 60bpm showing an increased amplitude at lower pacing on day 0 and a decreased amplitude at higher frequencies on day 4 in both groups, comprising 9-16 LMS generated by transverse slicing of embedded hearts. Functional testing was performed under electrical stimulation, with each stimulation frequency applied for a minimum of 10s under a preload of either 300 µN (top) or 1200 µN (bottom), targeting approximately 40 beats per frequency. TGF-β1 was applied as a profibrotic stimulus.

At early time points, LMS (both variants: LMS from isolated left ventricular tissue blocks and LMS prepared either by transverse slicing of embedded hearts) displayed pronounced frequency-dependent differences in contraction force, particularly at low stimulation frequencies (**Figure 7A-B**). With prolonged culture duration, this relationship progressively shifted, and contraction amplitude became less responsive to low frequencies and more dependent on higher pacing rates. Consistently, on day 0, contraction amplitude at 12 bpm was more than two-fold higher than the baseline measured at 60 bpm, whereas by day 4, stimulation at 240 bpm resulted in reduced amplitudes compared with the 60 bpm baseline in both preparation methods (**Figure 7A-B**). Taken together, these results demonstrate a clear, culture time-dependent alteration of the FFR in LMS. While these changes likely reflect modifications in tissue functionality and calcium handling, the underlying mechanisms driving this shift remain to be determined.

**Figure 7.**
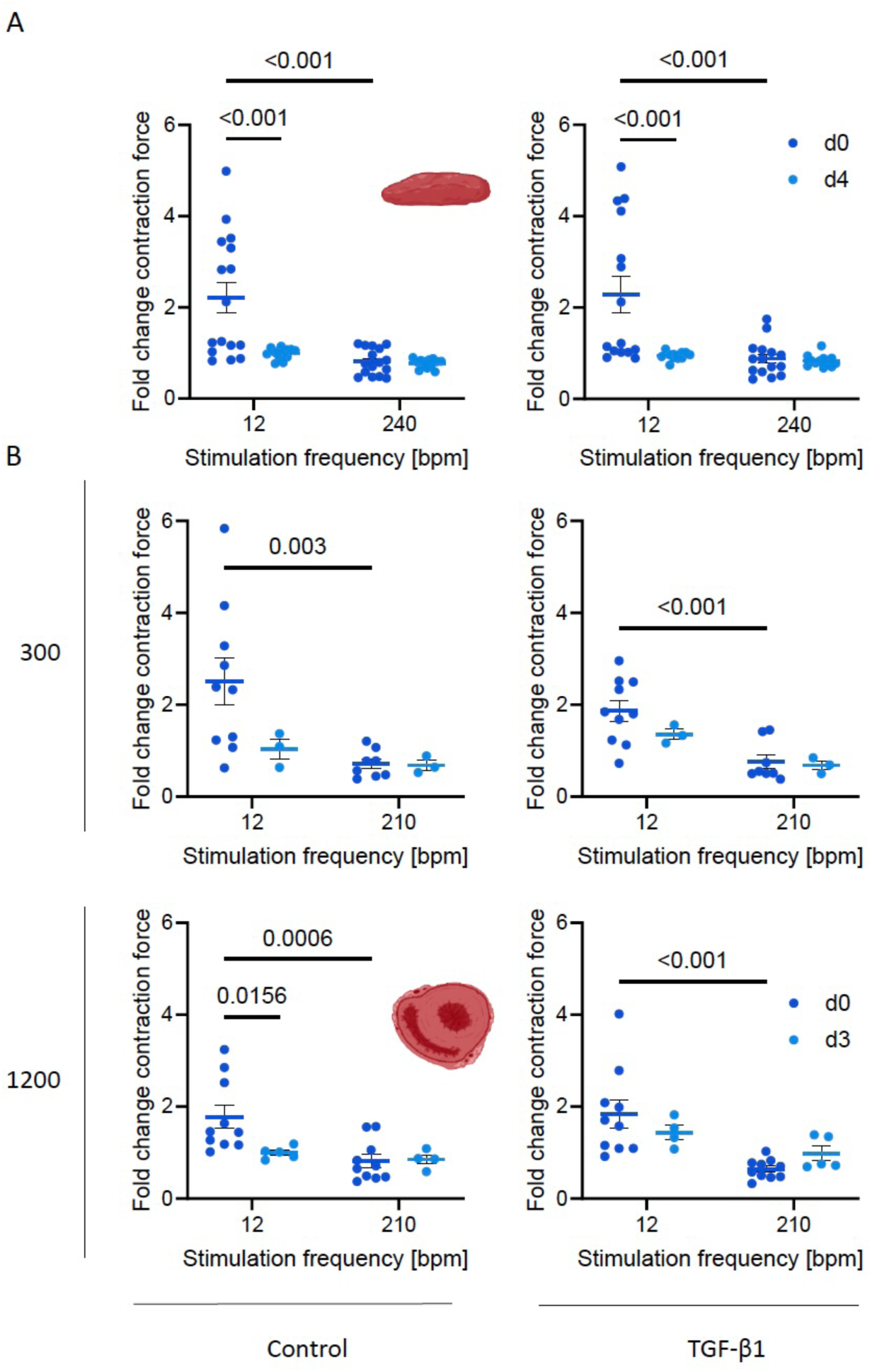
Force-frequency relationship of LMS changes with increasing pacing frequency. (**A**) Fold change of the contraction force shown at different frequencies of stimulation 12-210 bpm to the amplitude measured at 60 bpm (each sample to itself). LMS derived from isolated left ventricular tissue blocks. TGF-β1 was applied as a profibrotic stimulus. Pulled data from 4 experiments together leading to data from 6-16 samples. TGF-β1 was applied as a profibrotic stimulus. Two-way ANOVA, Uncorrected Fisher’s LSD, mean with SEM. (**B**) Fold changes calculated of individual sample to itself on amplitude measurements at 60bpm showing an increased amplitude at lower pacing on day 0 and a decreased amplitude at higher frequencies on day 4 in both groups, comprising 9-16 LMS generated by transverse slicing of embedded hearts. Functional testing was performed under electrical stimulation, with each stimulation frequency applied for a minimum of 10s under a preload of either 300 µN (top) or 1200 µN (bottom), targeting approximately 40 beats per frequency. TGF-β1 was applied as a profibrotic stimulus. Mixed effect model, Uncorrected Fisher’s LSD, mean with SEM. n= 14-16 biological replicates over 4 experiments.

The refractory period is classically defined as the interval following an action potential during which the initiation of a new action potential is unlikely or impossible. In the context of this study, we operationally define the refractory period as the time interval after a contraction during which a subsequent contraction cannot be elicited. To assess this, a predefined stimulation protocol was applied, in which the pacing frequency was gradually increased from 60 bpm to 300 bpm in 34 incremental steps, each maintained for 10 seconds, followed by 30-second recovery intervals at 60 bpm. Notably, even at the highest pacing rate of 300 bpm, two distinct contractions could still be evoked in the LMS. This behaviour was consistently observed throughout the entire culture period. Based on these findings, we conclude that, provided the tissue retained detectable contractile activity, the refractory period of mouse precision-cut myocardial tissue was shorter than 200 ms.

### 2.3. Induction and characterization of fibrosis in LMS

Collagen production is a key hallmark of fibrotic remodelling. To evaluate the effect of TGF-β1 stimulation on LMS across different culture conditions, pro-collagen I α1 (Pro-Col1α1) levels were quantified in culture supernatants by ELISA at multiple time points. Regardless of the culture method, pro-collagen I α1 secretion increased over time in both untreated and TGF-β1-treated LMS (**Figure 8A-C**). However, LMS maintained at the air-liquid interface secreted substantially lower levels of pro-collagen I α1 (**Figure 8A**) compared with LMS cultured in the biomimetic MyoDish system (**Figure 8B-C**). The highest secretion levels were observed in the biomimetic MyoDish system using LMS derived from isolated left ventricular tissue blocks (**Figure 8C**). Notably, pro-collagen I α1 secretion showed considerable inter-sample variability, likely reflecting differences in cell number, tissue origin within the heart, and overall tissue viability.

**Figure 8.**
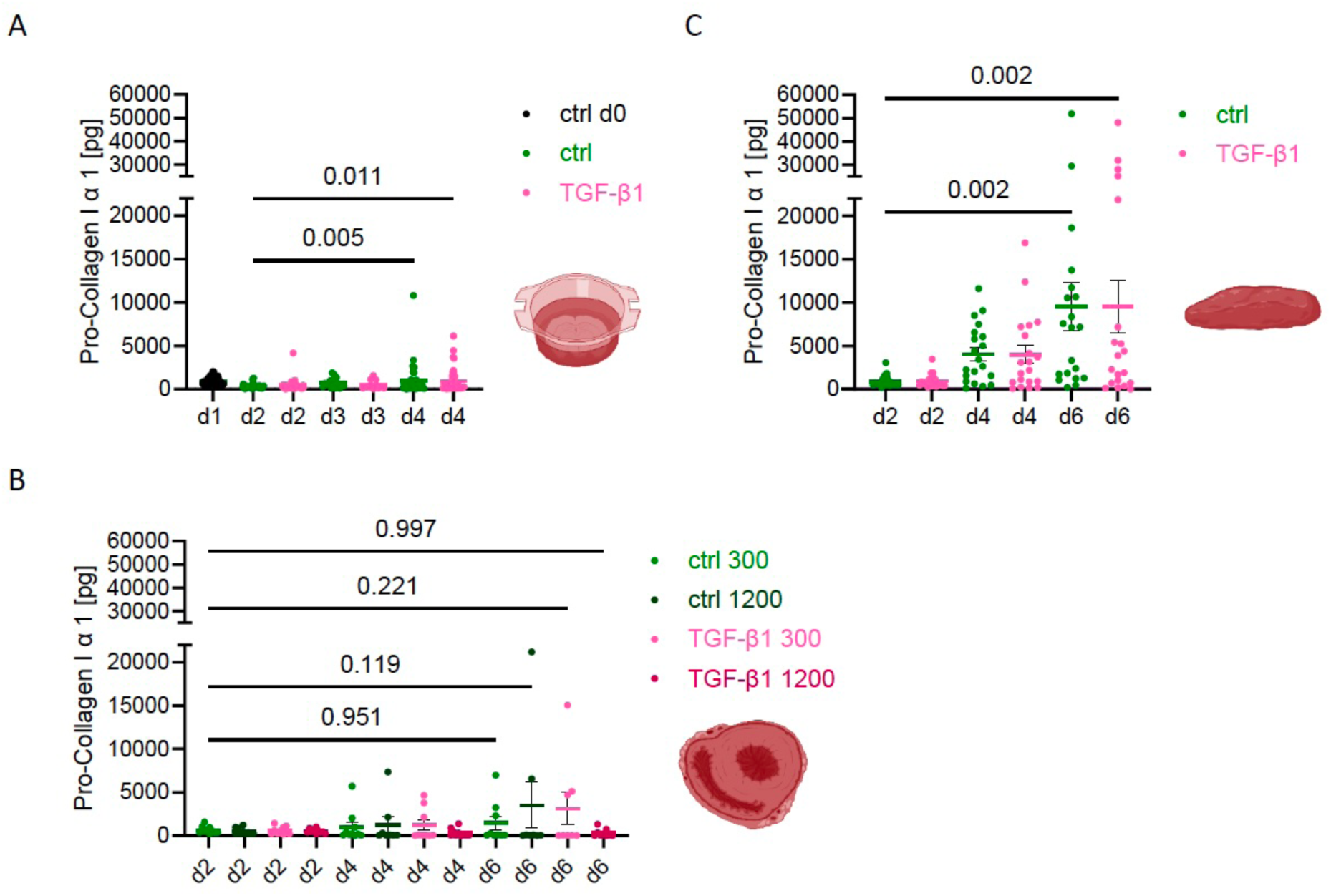
Pro-collagen I α1 secretion in LMS under fibrotic condition. Quantified comparison of stimulated (TGF-β1) and unstimulated (ctrl) condition during four days of LMS culture using the air-liquid interface system **(A**), during six days in the biomimetic MyoDish system with LMS obtained transverse slicing of embedded hearts using a pacing system (**B**) or isolated from left ventricular tissue blocks (**C**). n=36-75. Ordinary One-way ANOVA with Šídák’s multiple comparisons test. One data point represents one tissue cultured, meaning 16 biological replicates in 4 experiments. The sensitivity threshold of the assay is at 31.25 pg. Abbreviations: day 1 (d1), day 2 (d2), day 4 (d4), control/unstimulated group (ctrl), TGF-β1 (20 ng/mL) stimulated group (TGF-β1).

To further evaluate the culture methods and assess both structural integrity and fibrotic responses in LMS, histological and immunohistochemical analyses were performed. We focused on establishing robust staining protocols for hematoxylin and eosin (H&E) and Sirius Red to confirm tissue integrity and to visualize collagen deposition, respectively. In addition, immunostaining protocols were developed for key markers relevant to fibrotic remodelling, including von Willebrand factor, αSMA, podoplanin (gp38), vimentin, and periostin. **Figure 9** shows representative images of these stainings from LMS preserved on day 0 (unstimulated) and on day 4, with and without TGF-β1 stimulation. These analyses demonstrate that histological and immunohistochemical staining can be reliably performed on LMS and that overall tissue architecture remains well preserved both at baseline and after four days in culture under all tested conditions. TGF-β1 stimulation induced periostin expression, an indicator of activated fibroblasts, while reducing gp38 (podoplanin) expression associated with quiescent cells. Prolonged culture increased expression of both gp38 and vimentin (**Figure 9**).

**Figure 9.**
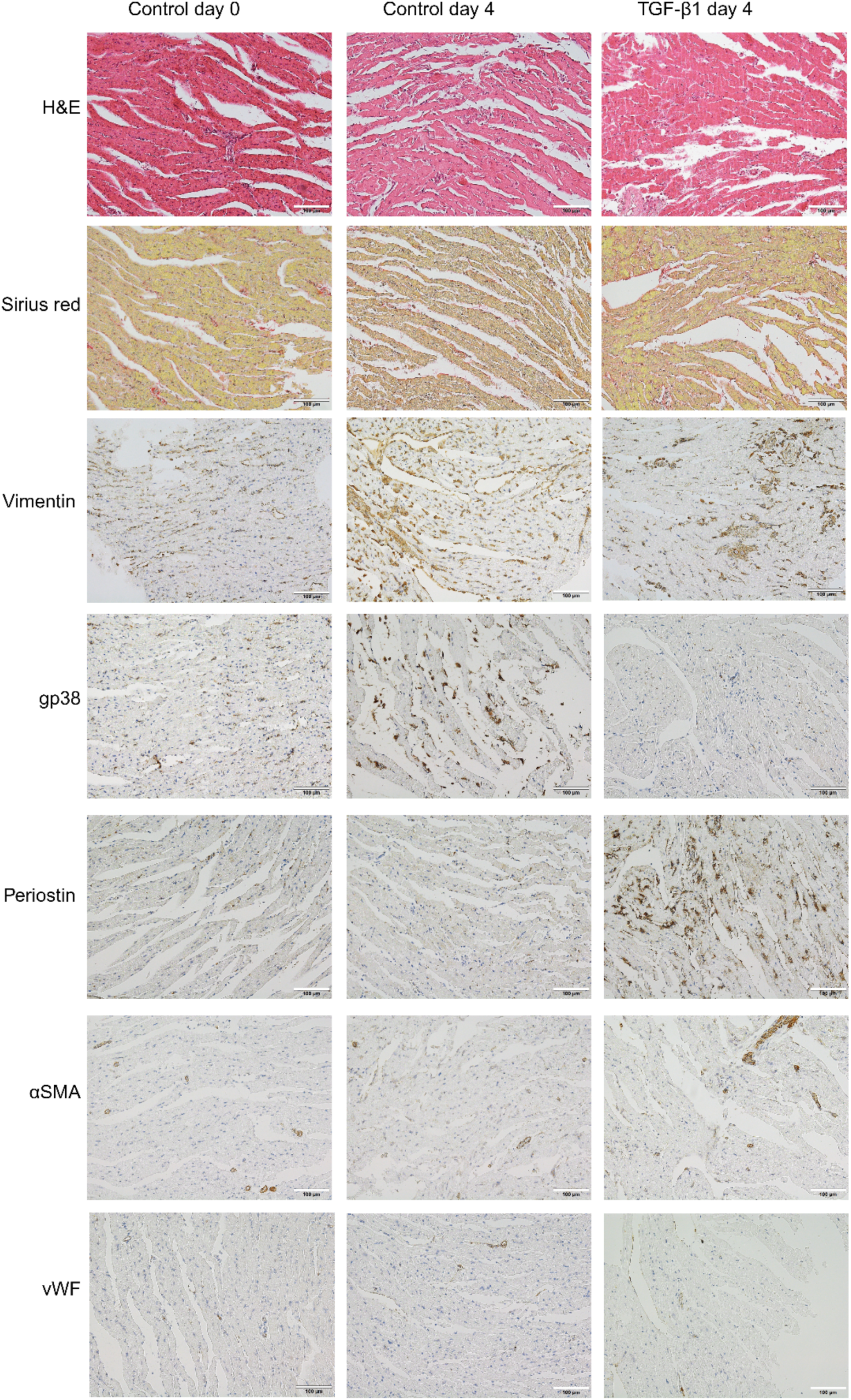
Representative histological and immunohistochemical images of LMS. Unstimulated LMS preserved on day 0 and LMS cultured under both conditions and preserved on day 4 are shown at 20× magnification. In immunohistochemical staining, nuclei were counterstained with hematoxylin. Scale bars: 100 µm. Abbreviations: α-smooth muscle actin (α-SMA), podoplanin (gp38), hematoxylin and eosin (H&E), von Willebrand factor (vWF). Bar = 100 μm.

## 3. Discussion

The development of experimental models that faithfully recapitulate the multicellular complexity, ECM composition, and biomechanical environment of the heart remains a critical challenge. An ideal in vitro model should reflect human myocardial biology, retain pathophysiological relevance, enable mechanistic insight, and support scalable experimentation^11^. 2D cultures of stem cell-derived or primary cardiac cells offer high experimental control, throughput, and suitability for long-term studies^11, 12^. However, they often lack multicellular complexity, native ECM interactions, and mature cellular phenotypes, limiting physiological relevance^10–13^. 3D cardiac models include whole-heart preparations, muscle-based systems, engineered tissues, and precision-cut LMS^10–13, 20, 21^. While whole-heart and muscle preparations closely resemble in vivo physiology, they are technically demanding, low throughput, and unsuitable for long-term culture. Engineered cardiac tissues provide controlled heterocellular environments but do not fully recapitulate native tissue architecture or mature functional properties. Precision-cut LMS combine advantages of 2D and 3D models by preserving native tissue structure, ECM, multicellularity, and mature contractility, while maintaining moderate experimental complexity and enabling longitudinal studies^10, 11, 13^. Compared with other 3D models, mouse LMS require fewer animals and allow cellular- and subcellular-level analyses. However, this approach is limited by the absence of vascular perfusion, substantial biological variability, and technical challenges associated with tissue preparation and analysis. A graphical overview of currently available in vitro and ex vivo cardiac models is presented in **Scheme 4**.

**Scheme 4.**
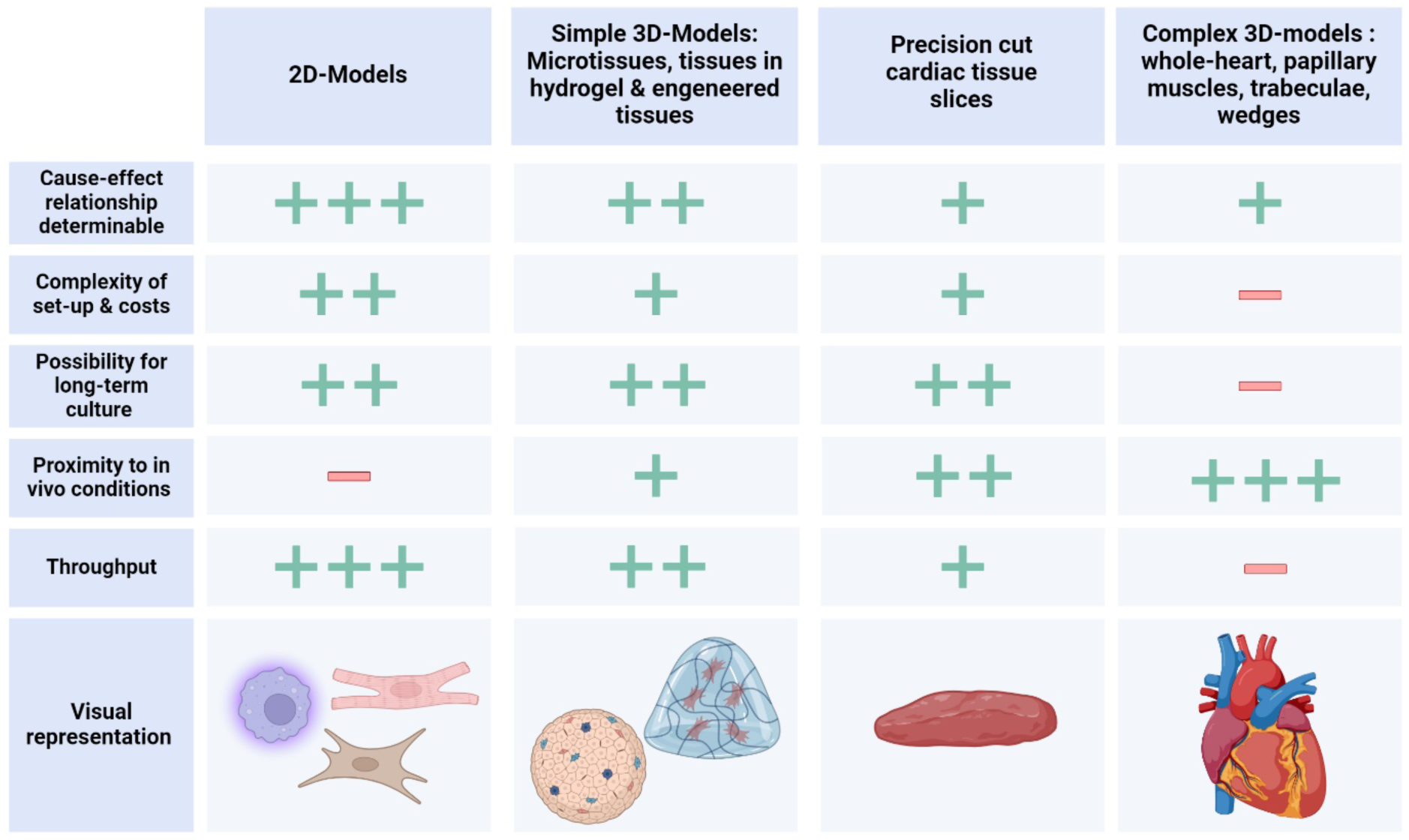
Comparison of cardiac research models. “+” and “–” indicate general advantages and limitations of each model, respectively. This figure was created using BioRender.com.

In this study, we established and evaluated murine LMS as a potential ex vivo model of cardiac fibrosis. We optimized vibratome-based tissue slicing and implemented two culture strategies: an air-liquid interface system and a biomimetic pacing-based MyoDish culture system. Tissue viability, contractile function, and molecular remodelling were systematically assessed, and the capacity of TGF-β1 stimulation to induce fibrotic remodelling in mouse LMS was evaluated. We demonstrated that LMS represent a promising intermediate model that bridges the gap between simplified 2D cultures and complex 3D systems, offering a valuable platform for mechanistic studies and drug testing in cardiac research. Together, our findings delineate the strengths and limitations of murine LMS for fibrosis research and identify critical parameters for further refinement of this model.

### Viability assessment of murine LMS

Assessing viability in thick myocardial tissue is inherently challenging and requires the integration of metabolic, visual, and functional readouts. To determine the maximal viable culture duration of murine LMS in our experimental setup, we applied several complementary approaches. Initial testing with ATP-based assays and PrestoBlue™ proved unreliable or insufficiently sensitive for myocardial tissue under our conditions (*data not shown*). We therefore employed the MTT assay as an alternative metabolic readout. While no marked decline in metabolic activity was detected over time in LMS cultured at the air-liquid interface, paced LMS in the biomimetic MyoDish showed a clear reduction over culture time, consistent with the observed decline in contractile function. Notably, LMS maintained in the biomimetic pacing MyoDish system displayed higher initial metabolic activity than those cultured at the air-liquid interface, indicating improved tissue preservation under biomimetic conditions. Interpretation of these findings is limited by the lack of normalization to tissue mass. Nevertheless, the MTT assay is well established in myocardial slice research and, when combined with complementary readouts, remains a useful indicator of tissue viability^12, 22–24^.

LIVE/DEAD staining provided additional spatial information and supported estimation of maximal culture duration. Using the LIVE/DEAD™ Viability/Cytotoxicity Kit, we observed increasing intra-slice heterogeneity over culture time, with visually distinct viable and non-viable regions and rising signal variability. By combining quantitative fluorescence analysis with visual inspection and comparison to dead controls, we concluded that murine LMS cultured at the air-liquid interface remain viable for up to four days. In paced LMS, however, tissue folding and increasing structural irregularity limited the applicability of staining-based approaches. Furthermore, even with confocal imaging, dye penetration is largely restricted to superficial cell layers, leaving deeper, and potentially more viable regions inaccessible^13, 25^. As a result, and in line with current trends in the field, functional and metabolic assessments are increasingly favoured over staining-based methods for evaluating viability in thick myocardial preparations.

### Functional assessment and implications for drug testing

As described above, the MyoDish pacing system enables continuous electrical and mechanical stimulation and allows real-time measurement of contraction amplitude. These readouts provide valuable information on tissue viability, functional integrity, and the potential development of pathological remodelling, including fibrosis. Importantly, electrical pacing and mechanical loading are essential to limit dedifferentiation and culture-induced remodelling of myocardial tissue^10^. Beyond force measurements, this model permits additional analyses, such as contraction kinetics^16, 26^, Ca²⁺ imaging^19^, protein expression studies, and histological or immunohistochemical evaluation. However, several technical and biological limitations must be considered, particularly when working with murine tissue. Owing to the relatively small size and lower contractile force of mouse myocardium, contraction amplitudes are small and the system is highly sensitive to movement of the spring wire. As a result, measurements are more susceptible to noise, for example from rocking of the sensor board, making functional analyses more challenging than in human or porcine slices, which generate larger forces. The following section therefore focuses on the relevance of force-frequency relationship (FFR) and post-rest potentiation (PRP) measurements, as well as the influence of tissue origin, for functional assessment and drug testing in LMS.

PRP reflects Ca²⁺ accumulation within the sarcoplasmic reticulum during pauses in stimulation and provides insight into the tissue’s capacity to store and release Ca²⁺. Similarly, the force–frequency relationship is closely linked to intracellular Ca²⁺ handling, with a negative FFR being associated with altered Ca²⁺ homeostasis and reduced T-tubular density^19^. Thus, analysis of PRP and FFR enables indirect assessment of excitation–contraction coupling and overall tissue functionality. These functional parameters are particularly relevant for drug testing, as cardiotoxicity remains a leading cause of drug withdrawal and adverse drug reactions frequently involve impaired contractility, arrhythmias, or heart failure^26^. In heart failure, contractile dysfunction is commonly linked to disturbances in Ca²⁺ homeostasis^27^. Accordingly, incorporating Ca²⁺-related functional readouts such as PRP and FFR into pharmacological studies may improve both drug screening and safety assessment^28^. Previous studies using precision-cut LMS, primarily from larger species, have demonstrated the high translational relevance of these approaches^19, 20, 26^.

Physiologically, cardiac tissue exhibits a positive FFR, which can become negative in chronic heart failure^29^. In fresh porcine and human myocardial slices, a positive FFR has been reported, with a gradual inversion during prolonged culture^26^. This shift has been linked to T-tubule remodelling and altered sarcoplasmic reticulum Ca²⁺ handling^26, 29–31^. In our study, PRP was clearly detectable on day 0 but rapidly diminished with time in culture. Similarly, the FFR evolved over time: on day 0, contraction amplitude was increased at lower pacing frequencies, while higher frequencies did not significantly alter force compared with the baseline of 60 bpm. By day 4, this relationship was inverted, with lower frequencies having little effect and higher frequencies causing a reduction in contraction amplitude. These findings indicate a progressive decline in contractile functionality during culture. Potential explanations include disturbed Ca²⁺ homeostasis, for example due to altered expression or function of SERCA or NCX proteins involved in cytosolic Ca²⁺ removal^19, 26^, reduced tissue viability with fewer actively contracting cells, or culture-induced tissue remodelling such as changes in extracellular matrix composition leading to increased stiffness and impaired force generation.

Overall, our data suggest that although murine precision-cut LMS remain viable and contractile for up to four days, functional performance declines rapidly. This substantially limits the time window for pharmacological studies, particularly for compounds targeting Ca²⁺ signalling pathways, which in our current setup can likely only be reliably assessed within the first 48 hours of culture.

### TGF-β1 stimulation to induce fibrosis in murine LMS

A central aim of this study was to establish a murine LMS-based model of cardiac fibrosis. To evaluate whether TGF-β1 stimulation can induce fibrosis in LMS, we combined metabolic, functional, and molecular readouts. The MTT viability assay revealed a significant difference in metabolic activity between TGF-β1-stimulated and control LMS. This observation could indicate an enhanced activation of apoptotic pathways downstream of TGF-β1 receptor type II signalling [62,63]. However, several methodological limitations must be considered. Tissue mass was not normalized between samples, and given the slicing procedure, control samples may on average have contained more tissue than TGF-β1-treated samples. Because the MTT assay reflects overall metabolic activity, differences in tissue amount could bias the results.

Consistent with this interpretation, RT-qPCR, ELISA, and contraction analyses did not reveal robust or reproducible differences between control and TGF-β1-stimulated LMS. We therefore conclude that, under our experimental conditions, TGF-β1 stimulation at 20 ng/mL is insufficient to reliably induce fibrosis in mouse LMS. Potential explanations include activation of counter-regulatory or apoptotic pathways via TGF-β1 receptor signalling [62,63], insufficient concentration or duration of stimulation, or strong baseline remodelling induced by the culture conditions themselves. Indeed, the pronounced increase in pro-collagen 1α1 secretion observed in paced slices suggests that the biomimetic MyoDish culture environment may already promote substantial tissue remodelling or fibrotic signalling^10^, potentially masking additional effects of exogenous TGF-β1. Notably, TGF-β1 at the same concentration robustly induces fibroblast activation and fibrosis in 3D cardiac microtissues composed of human iPSC-derived cardiomyocytes and cardiac fibroblasts^21, 32^. The divergent responses between these models likely reflect fundamental differences in cellular composition, ECM presence and remodelling, tissue density, and endogenous regulatory mechanisms. In contrast to simplified microtissues, the complex multicellular environment of LMS may allow for buffering or compensatory responses that limit fibrotic remodelling.

To establish a robust fibrosis model using LMS, several alternative strategies could be considered. Increasing the concentration of TGF-β1 may enhance fibrotic signalling but would further deviate from physiological levels^33–35^ and increase the risk of toxicity and apoptosis. Alternatively, TGF-β1 could be combined with additional profibrotic stimuli, such as IL-11^20^, although this may complicate interpretation of cause-effect relationships. Finally, fibrosis could be induced using different stimuli, including TNF-α, IL-1, or IL-11^4^, or by applying increased mechanical preload to the tissue^11, 20, 36^, thereby more closely mimicking disease-relevant biomechanical stress.

Analysis of procollagen 1α1 secretion provided additional insight into tissue viability and remodelling. In LMS cultured within the pacing MyoDish system, procollagen 1α1 levels increased over time, whereas slices maintained at the air-liquid interface showed a decline beginning on day 2. Reduced procollagen 1α1 secretion in air-liquid interface cultures likely reflects early functional deterioration and loss of viable fibroblasts. In contrast, increased procollagen 1α1 production under paced conditions may indicate enhanced tissue viability but may also reflect culture-induced remodelling in response to altered mechanical and metabolic demands. Mechanical load and electrical stimulation are known to preserve cardiomyocyte differentiation and functionality while reducing adverse remodelling^10^, supporting our conclusion that biomimetic MyoDish culture conditions are superior to static air-liquid interface culture. Functional measurements further supported these findings. Contraction amplitude, serving as a proxy for force generation, stabilized after an initial adaptation period of approximately 24 h before declining after day 4. In parallel, stimulation threshold increased over time, indicating progressive impairment of excitation-contraction coupling and tissue responsiveness. Together, contraction amplitude and stimulation threshold measurements corroborated the metabolic viability data and suggested that murine LMS cultured in a pacing MyoDish system maintain stable viability and functionality for approximately 4 days.

Across all assays, substantial variability was observed between hearts and between LMS from the same heart. This highlights an inherent challenge of LMS models and underscores the importance of combining multiple viability and functional readouts, particularly during model establishment. While functional measurements are non-terminal and directly reflect cardiac performance, they should ideally be interpreted in conjunction with metabolic and molecular assessments to account for biological variability.

### Advantages and limitations of murine LMS

Murine LMS offer advantages such as accessibility, cost efficiency, and compatibility with genetic disease models. However, our findings indicate that many benefits of the LMS approach are diminished when using mouse tissue. The small heart size limits slice yield, inter-sample variability is high, and rapid metabolism complicates physiological pacing and functional assessment. Moreover, the geometry of the mouse myocardium hinders isolation of discrete fiber sheets, which may contribute to reduced viability^37^. These limitations contrast with LMS derived from larger mammals or human tissue, which show superior long-term viability and functional stability^37–39^.

### Conclusions

In summary, we established and characterized murine LMS and compared air-liquid interface and biomimetic MyoDish pacing-system culture conditions. We demonstrate that MyoDish pacing-system culture preserves viability and functionality more effectively, supporting stable LMS culture for up to 6 days. Under experimental conditions, TGF-β1 stimulation alone was insufficient to induce robust fibrosis in murine LMS. While mouse LMS offer certain advantages, their limitations suggest that long-term fibrosis modelling and pharmacological studies may be better suited to LMS derived from larger animals or human tissue. Nevertheless, LMS remain a powerful translational platform, and continued refinement of culture conditions and stimulation paradigms will further enhance their value in cardiac fibrosis research.

## Acknowledgment

We acknowledge the LASC animal facility in Schlieren, UZH, for their excellent animal care and support.

## Funding

This work was supported by the Swiss National Science Foundations (310030_175663, 310030_20770).

## Conflict of interest statement (in addition to author disclosure form)

The disclosure statement of Prof. O. Distler (3 years backwards, 2024-2026): Prof. O. Distler has/had consultancy relationship with and/or has received research funding from or has served as a speaker for the following companies in the area of potential treatments for systemic sclerosis and its complications in the last two years: 4P-Pharma, Abbvie, Acceleron, Alcimed, Altavant, Amgen, AnaMar, Arxx, AstraZeneca, Blade, Boehringer Ingelheim, Citus AG, Corbus, CSL Behring, Galderma, Galapagos, Glenmark, Gossamer, Kymera, Lupin, Medscape, Merck, Miltenyi Biotec, Mitsubishi Tanabe; Novartis, Prometheus, Redxpharna, Roivant and Topadur in the area of potential treatments of scleroderma and its complications. Patent issued “mir-29 for the treatment of systemic sclerosis” (US8247389, EP2331143). Co-founder of CITUS. Research Grants: Kymera, Mitsubishi Tanabe. Prof. R. Lafyatis reports grants from Bristol Meyer Squib, Corbus, Formation, Moderna, Regeneron Bain Capital and Pfizer, outside the submitted work; served or serves as a consultant with Bristol Meyers Squibb, Formation, Thirona Bio, Sanofi, Boehringer-Ingelheim, Merck, Genentech/Roche, EMD Serono, Morphic, Third Rock Ventures and Zag Bio; and sits on an independent data safety monitoring committees for Advarra/GSK and Genentech. All other authors declare no competing interests.

## Notes

### Competing Interest Statement

The authors have declared no competing interest.

## References

1. Vaduganathan M, Mensah GA, Turco JV, Fuster V, Roth GA. The Global Burden of Cardiovascular Diseases and Risk: A Compass for Future Health. J Am Coll Cardiol. 2022;80:2361–2371.

2. Rabe E. Cyanoacrylate adhesive in the treatment of saphenous vein incompetence. Comment on Calik et al, page 241-246. *Vasa*. 2016;45:193–194.

3. Global Burden of Cardiovascular D, Risks C. Global, Regional, and National Burden of Cardiovascular Diseases and Risk Factors in 204 Countries and Territories, 1990-2023. J Am Coll Cardiol. 2025;86:2167–2243.

4. Frangogiannis NG. Cardiac fibrosis. Cardiovasc Res. 2021;117:1450–1488.

5. Maruyama K, Imanaka-Yoshida K. The Pathogenesis of Cardiac Fibrosis: A Review of Recent Progress. Int J Mol Sci. 2022;23.

6. Wong TC, Piehler K, Meier CG, Testa SM, Klock AM, Aneizi AA et al. Association between extracellular matrix expansion quantified by cardiovascular magnetic resonance and short-term mortality. Circulation. 2012;126:1206–1216.

7. Kong P, Christia P, Frangogiannis NG. The pathogenesis of cardiac fibrosis. Cell Mol Life Sci. 2014;71:549–574.

8. Pinto AR, Ilinykh A, Ivey MJ, Kuwabara JT, D’Antoni ML, Debuque R et al. Revisiting Cardiac Cellular Composition. Circ Res. 2016;118:400–409.

9. Jugdutt BI. Ventricular remodeling after infarction and the extracellular collagen matrix: when is enough enough? Circulation. 2003;108:1395–1403.

10. Watson SA, Dendorfer A, Thum T, Perbellini F. A practical guide for investigating cardiac physiology using living myocardial slices. Basic Res Cardiol. 2020;115:61.

11. Pitoulis FG, Watson SA, Perbellini F, Terracciano CM. Myocardial slices come to age: an intermediate complexity in vitro cardiac model for translational research. Cardiovasc Res. 2020;116:1275–1287.

12. Miller JM, Meki MH, Elnakib A, Ou Q, Abouleisa RRE, Tang XL et al. Biomimetic cardiac tissue culture model (CTCM) to emulate cardiac physiology and pathophysiology ex vivo. Commun Biol. 2022;5:934.

13. Watson SA, Scigliano M, Bardi I, Ascione R, Terracciano CM, Perbellini F. Preparation of viable adult ventricular myocardial slices from large and small mammals. Nat Protoc. 2017;12:2623–2639.

14. Watson SA, Scigliano M, Bardi I, Ascione R, Terracciano CM, Perbellini F. Preparation of viable adult ventricular myocardial slices from large and small mammals. Nature Protocols. 2017;12:2623–2639.

15. Cao-Ehlker X, Fischer C, Lu K, Bruegmann T, Sasse P, Dendorfer A et al. Optimized Conditions for the Long-Term Maintenance of Precision-Cut Murine Myocardium in Biomimetic Tissue Culture. Bioengineering (Basel). 2023;10.

16. Hamers J, Sen P, Merkus D, Seidel T, Lu K, Dendorfer A. Preparation of Human Myocardial Tissue for Long-Term Cultivation. J Vis Exp. 2022.

17. Fischer C, Milting H, Fein E, Reiser E, Lu K, Seidel T et al. Long-term functional and structural preservation of precision-cut human myocardium under continuous electromechanical stimulation in vitro. Nature Communications. 2019;10.

18. Livak KJ, Schmittgen TD. Analysis of Relative Gene Expression Data Using Real-Time Quantitative PCR and the 2−ΔΔCT Method. Methods. 2001;25:402–408.

19. Sun Z, Lu K, Kamla C, Kameritsch P, Seidel T, Dendorfer A. Synchronous force and Ca(2+) measurements for repeated characterization of excitation-contraction coupling in human myocardium. Commun Biol. 2024;7:220.

20. Nunez-Toldra R, Kirwin T, Ferraro E, Pitoulis FG, Nicastro L, Bardi I et al. Mechanosensitive molecular mechanisms of myocardial fibrosis in living myocardial slices. ESC Heart Fail. 2022;9:1400–1412.

21. Blyszczuk P, Zuppinger C, Costa A, Nurzynska D, Di Meglio FD, Stellato M et al. Activated Cardiac Fibroblasts Control Contraction of Human Fibrotic Cardiac Microtissues by a beta-Adrenoreceptor-Dependent Mechanism. Cells. 2020;9.

22. Ross AJ, Krumova I, Tunc B, Wu Q, Wu C, Camelliti P. A novel method to extend viability and functionality of living heart slices. Front Cardiovasc Med. 2023;10:1244630.

23. Klumm MJ, Heim C, Fiegle DJ, Weyand M, Volk T, Seidel T. Long-Term Cultivation of Human Atrial Myocardium. Front Physiol. 2022;13:839139.

24. Pfeuffer AM, Kupfer LK, Shankar TS, Drakos SG, Volk T, Seidel T. Ryanodine Receptor Staining Identifies Viable Cardiomyocytes in Human and Rabbit Cardiac Tissue Slices. Int J Mol Sci. 2023;24.

25. Perbellini F, Watson SA, Scigliano M, Alayoubi S, Tkach S, Bardi I et al. Investigation of cardiac fibroblasts using myocardial slices. Cardiovasc Res. 2018;114:77–89.

26. Shi R, Reichardt M, Fiegle DJ, Kupfer LK, Czajka T, Sun Z et al. Contractility measurements for cardiotoxicity screening with ventricular myocardial slices of pigs. Cardiovasc Res. 2023;119:2469–2481.

27. Pieske B, Maier LS, Schmidt-Schweda S. Sarcoplasmic reticulum Ca2+ load in human heart failure. Basic Res Cardiol. 2002;97 Suppl 1:I63–71.

28. Fischer C, Milting H, Fein E, Reiser E, Lu K, Seidel T et al. Long-term functional and structural preservation of precision-cut human myocardium under continuous electromechanical stimulation in vitro. Nat Commun. 2019;10:117.

29. Abu-Khousa M, Fiegle DJ, Sommer ST, Minabari G, Milting H, Heim C et al. The Degree of t-System Remodeling Predicts Negative Force-Frequency Relationship and Prolonged Relaxation Time in Failing Human Myocardium. Front Physiol. 2020;11:182.

30. Alpert NR, Leavitt BJ, Ittleman FP, Hasenfuss G, Pieske B, Mulieri LA. A mechanistic analysis of the force-frequency relation in non-failing and progressively failing human myocardium. Basic Res Cardiol. 1998;93 Suppl 1:23–32.

31. Mulieri LA, Hasenfuss G, Leavitt B, Allen PD, Alpert NR. Altered myocardial force-frequency relation in human heart failure. Circulation. 1992;85:1743–1750.

32. Lee MO, Jung KB, Jo SJ, Hyun SA, Moon KS, Seo JW et al. Modelling cardiac fibrosis using three-dimensional cardiac microtissues derived from human embryonic stem cells. J Biol Eng. 2019;13:15.

33. Khan SA, Joyce J, Tsuda T. Quantification of active and total transforming growth factor-beta levels in serum and solid organ tissues by bioassay. BMC Res Notes. 2012;5:636.

34. Gomez-Bernal F, Quevedo-Abeledo JC, Garcia-Gonzalez M, Fernandez-Cladera Y, Gonzalez-Rivero AF, de Vera-Gonzalez A et al. Serum Levels of Transforming Growth Factor Beta 1 in Systemic Lupus Erythematosus Patients. Biomolecules. 2022;13.

35. Wakefield LM, Letterio JJ, Chen T, Danielpour D, Allison RS, Pai LH et al. Transforming growth factor-beta1 circulates in normal human plasma and is unchanged in advanced metastatic breast cancer. Clin Cancer Res. 1995;1:129–136.

36. Watson SA, Duff J, Bardi I, Zabielska M, Atanur SS, Jabbour RJ et al. Biomimetic electromechanical stimulation to maintain adult myocardial slices in vitro. Nat Commun. 2019;10:2168.

37. Plikus MV, Wang X, Sinha S, Forte E, Thompson SM, Herzog EL et al. Fibroblasts: Origins, definitions, and functions in health and disease. Cell. 2021;184:3852–3872.

38. Shinde AV, Humeres C, Frangogiannis NG. The role of alpha-smooth muscle actin in fibroblast-mediated matrix contraction and remodeling. Biochim Biophys Acta Mol Basis Dis. 2017;1863:298–309.

39. Tomasek JJ, Gabbiani G, Hinz B, Chaponnier C, Brown RA. Myofibroblasts and mechano-regulation of connective tissue remodelling. Nat Rev Mol Cell Biol. 2002;3:349–363.

